# Coupling of Slack and Na_V_1.6 sensitizes Slack to quinidine blockade and guides anti-seizure strategy development

**DOI:** 10.1101/2023.03.16.532982

**Authors:** Tian Yuan, Yifan Wang, Yuchen Jin, Hui Yang, Shuai Xu, Heng Zhang, Qian Chen, Na Li, Xinyue Ma, Huifang Song, Chao Peng, Ze Geng, Jie Dong, Guifang Duan, Qi Sun, Yang Yang, Fan Yang, Zhuo Huang

**Affiliations:** State Key Laboratory of Natural and Biomimetic Drugs, Department of Molecular and Cellular Pharmacology, School of Pharmaceutical Sciences, Peking University Health Science Center, Beijing, 100191, China; Department of Medicinal Chemistry and Molecular Pharmacology, College of Pharmacy, Purdue University, _West Lafayette_, IN 47907, USA; Department of Biophysics, Kidney Disease Center of the First Affiliated Hospital, Zhejiang University School of Medicine, Hangzhou, _Zhejiang_, 310058, China; NHC and CAMS Key Laboratory of Medical Neurobiology, MOE Frontier Science Center for Brain Research and Brain-Machine Integration, School of Brain Science and Brain Medicine, Zhejiang University, Hangzhou, _Zhejiang_, 310058, China; IDG/McGovern Institute for Brain Research, Peking University, Beijing, 100871, China

**Keywords:** Slack, Na_V_1.6, quinidine, KCNT1-related epilepsy

## Abstract

Quinidine has been used as an anticonvulsant to treat patients with KCNT1-related epilepsy by targeting gain-of-function KCNT1 pathogenic mutant variants. However, the detailed mechanism underlying quinidine’s blockade against KCNT1 (Slack) remains elusive. Here, we report a functional and physical coupling of the voltage-gated sodium channel Na_V_1.6 and Slack. Na_V_1.6 binds to and highly sensitizes Slack to quinidine blockade. Homozygous knockout of Na_V_1.6 reduces the sensitivity of native sodium-activated potassium currents to quinidine blockade. Na_V_1.6-mediated sensitization requires the involvement of Na_V_1.6’s N-and C-termini binding to Slack’s C-terminus, and is enhanced by transient sodium influx through Na_V_1.6. Moreover, disrupting the Slack-Na_V_1.6 interaction by viral expression of Slack’s C-terminus can protect against SlackG269S-induced seizures in mice. These insights about a Slack-Na_V_1.6 complex challenge the traditional view of “Slack as an isolated target” for anti-epileptic drug discovery efforts, and can guide the development of innovative therapeutic strategies for KCNT1-related epilepsy.

**GRAPHICAL ABSTRACT:** 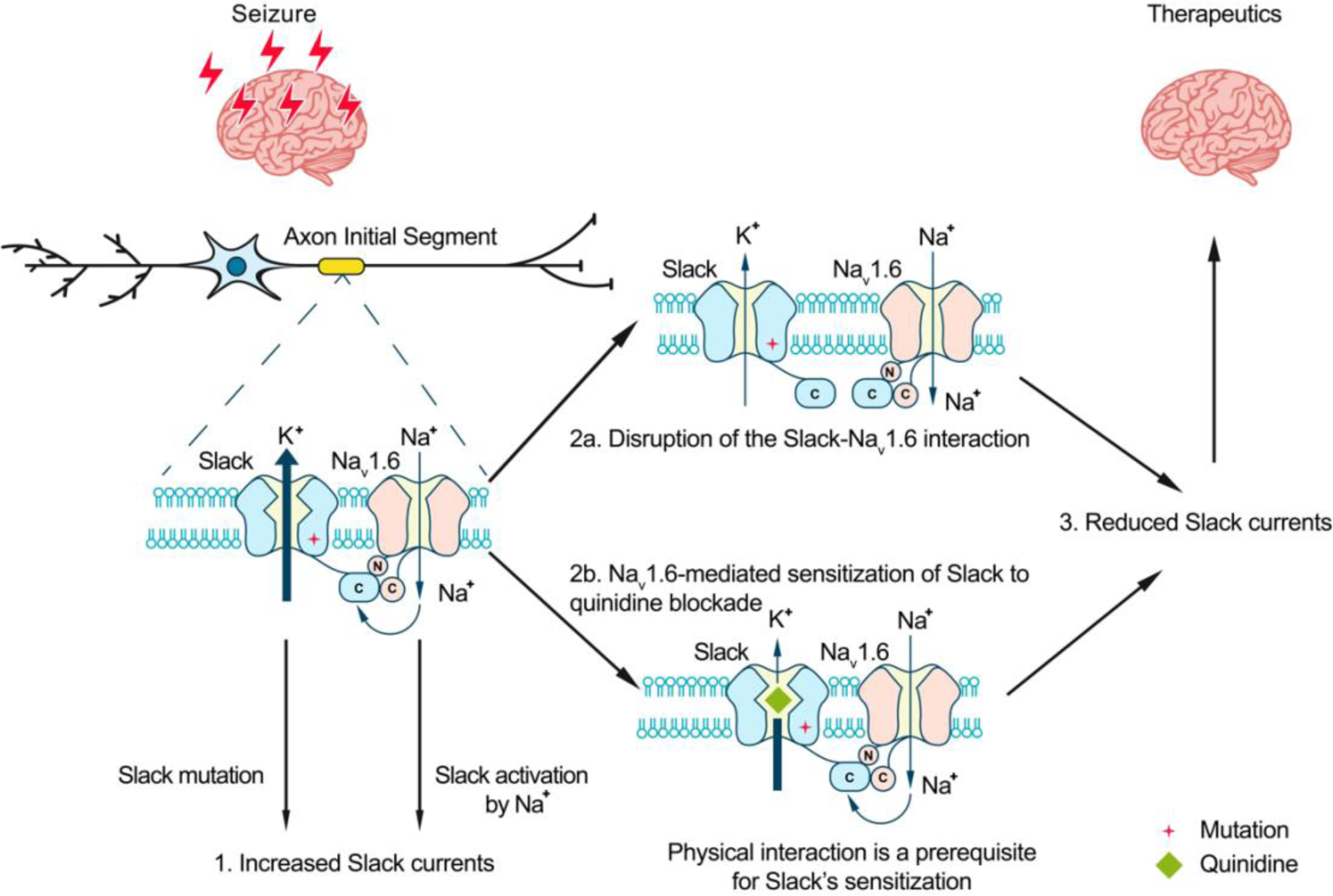

## INTRODUCTION

The sodium-activated potassium (K_Na_) channels Slack and Slick were first identified in guinea pig cardiomyocytes, and were subsequently found to be encoded by two genes of the *Slo2* family^1,2^. Slack channels encoded by the *Slo2.2* (*KCNT1*) gene are gated by Na^+^, while Slick channels encoded by the *Slo2.1* (*KCNT2*) gene are more sensitive to Cl^-^ than Na^+^. Slack channels are expressed at high levels in the central nervous system (CNS), especially the cortex and brainstem^3–5^. Activation of Slack channels by intracellular sodium ions forms delayed outward currents in neurons, contributes to slow afterhyperpolarization (AHP) following repeated action potentials, and modulates the firing frequency of neurons^6,7^.

Mutations in the *KCNT1* gene have been implicated in a wide spectrum of epileptic disorders, including early-onset epilepsy (*e.g.*, epilepsy of infancy with migrating focal seizures (EIMFS), non-EIMFS developmental and epileptic encephalopathies, and autosomal dominant or sporadic sleep-related hypermotor epilepsy (ADSHE))^8–11^. Over 50 mutations related to seizure disorders have been identified, typically displaying a gain-of-function (GOF) phenotype in heterologous expression systems^11,12^. The prescribed antiarrhythmic drug, quinidine, has emerged as a precision therapy for KCNT1-related epilepsy by blocking Slack mutant variants *in vitro* and conferring decreased seizure frequency and improved psychomotor development in clinical treatment^13–16^. However, clinical quinidine therapy has shown limited success and contradictory therapeutic effects, probably due to poor blood–brain barrier penetration, dose-limiting off-target effects, phenotype-genotype associations, and rational therapeutic schedule^16–19^.

Slack requires high intracellular free Na^+^ concentrations ([Na^+^]_in_) for its activation in neurons (K_d_ of ∼66 mM)^20^. However, previous investigations have shown that the [Na^+^]_in_ at resting states (∼10 mM) is much lower than the [Na^+^]_in_ needed for effective Slack activation (*e.g.*, K_d_ value)^1,20,21^. Therefore, native Slack channels need to localize with Na^+^ sources within a nanodomain to be activated and exert physiological function. Slack channels are known to be functionally coupled with sodium-permeable ion channels in neurons, such as voltage-gated sodium (Na_V_) channels and AMPA receptors ^21,22^. A question arises as to whether these known Na^+^ sources modulate Slack’s sensitivity to quinidine blockade.

Here, we found that Na_V_1.6 sensitizes Slack to quinidine blockade. Slack and Na_V_1.6 form a complex that functions in Na_V_1.6-mediated transient sodium influx to sensitize Slack to quinidine blockade in HEK293 cells and in primary cortical neurons. The widespread expression of these channel proteins in cerebral cortex, hippocampus, and cerebellum supports that the Na_V_1.6-Slack complex is essential for the function of a wide range of electrically excitable neurons, and moreover, that this complex can be viewed as a vulnerable target for drug development to treat KCNT1-related disorders.

## RESULTS

### Na_V_1.6 sensitizes Slack to quinidine blockade

Slack currents are activated by sodium entry through voltage-gated sodium (Na_V_) channels and ionotropic glutamate receptors (*e.g.* AMPA receptors)^21–23^. To investigate potential modulators of Slack’s sensitivity to quinidine blockade, we initially focused on the known Na^+^ sources of Slack. Working in HEK293 cells, we co-expressed Slack with AMPA receptor subunits (GluA1, GluA2, GluA3, or GluA4) or Na_V_ channel α subunits (Na_V_1.1, Na_V_1.2, Na_V_1.3, or Na_V_1.6), which are highly expressed in the central nervous system^24–27^. The sensitivity of Slack to quinidine blockade was assessed based on the detected inhibitory effects of quinidine on delayed outward potassium currents^13,28^. Interestingly, all neuronal Na_V_ channels significantly sensitized Slack to 30 μM quinidine blockade, whereas no effect was observed upon co-expression of Slack with GluA1, GluA2, GluA3, or GluA4 (Fig. S1).

When co-expressing Slack with Na_V_1.6 in HEK293 cells, Na_V_1.6 sensitized Slack to quinidine blockade by nearly 100-fold (IC_50_ = 85.13 μM for Slack expressed alone and an IC_50_ = 0.87 μM for Slack upon co-expression with Na_V_1.6) (Fig.1A,B,D,F). Na_V_1.6 also exhibited >10-fold selectivity in sensitizing Slack to quinidine blockade against Na_V_1.1, Na_V_1.2, and Na_V_1.3 (Fig. 1C,F, and Supplementary Table 1). When we co-expressed the cardiac sodium channel Na_V_1.5 with Slack, we observed only ∼3-fold sensitization of Slack to quinidine blockade (Fig.1E,F). These results together indicate that the apparent functional coupling between Slack and Na_V_ channels is Na_V_-channel-subtype-specific, with Na_V_1.6 being particularly impactful in Slack’s responsivity to quinidine blockade.

**Figure 1.**
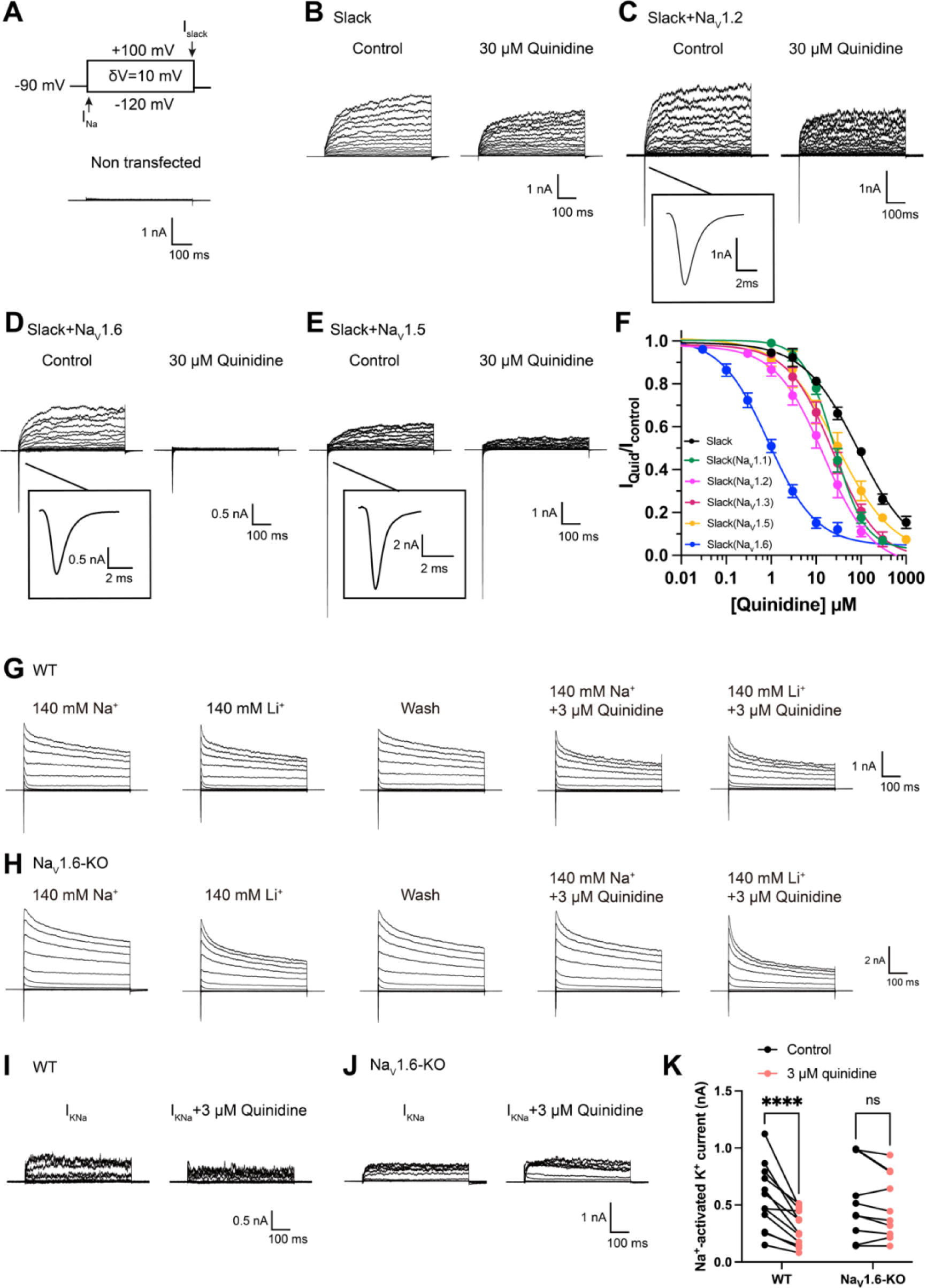
Na_V_1.6 specifically sensitizes Slack to quinidine blockade. (**A**) The voltage protocol and current traces from control (non-transfected) HEK293 cells. The arrows on the voltage protocol indicate the onset of inward sodium currents through Na_V_ channels and delayed outward potassium currents through Slack channels. The currents were evoked by applying 600-ms step pulses to voltages varying from -120 mV to +100 mV in 10 mV increments, with a holding potential of -90 mV and a stimulus frequency of 0.20 Hz. (**B**) Example current traces from HEK293 cells expressing Slack alone. The left traces show the family of control currents; the right traces show Slack currents remaining after application of 30 μΜ quinidine in the bath solution. (**C-E**) Example current traces from HEK293 cells co-expressing Slack with Na_V_1.2 (**C**), Na_V_1.6 (**D**), or Na_V_1.5 (**E**) channels before and after application of 30 μΜ quinidine. (**F**) The concentration-response curves for blocking of Slack by quinidine at +100 mV upon expression of Slack alone (n = 6) and co-expression of Slack with Na_V_1.1 (n = 7), Na_V_1.2 (n = 10), Na_V_1.3 (n = 13), Na_V_1.5 (n = 9), or Na_V_1.6 (n = 19). Please refer to Supplementary Table 1 for IC_50_ values. (**G-H**) Delayed outward currents in primary cortical neurons from postnatal (P0-P1) homozygous Na_V_1.6 knockout C3HeB/FeJ mice (Na_V_1.6-KO) (**H**) and the wild-type littermate controls (WT) (**G**). Current traces were elicited by 600-ms step pulses to voltages varying from -120 mV to +100 mV in 20 mV increments, with a holding potential of -70 mV, and recorded with different bath solutions in the following order: Na^+^-based bath solution (I_Control_), replacement of external Na^+^ with Li^+^ in equivalent concentration (I_Li_), washout of quinidine by Na^+^-based bath solution (I_Wash_), Na^+^-based bath solution with 3 μM quinidine (I_Quid_), Li^+^-based bath solution with 3 μM quinidine (I_Li+Quid_). The removal and subsequent replacement of extracellular Na^+^ revealed the I_KNa_ in neurons. (**I-J**), The sensitivity of native sodium-activated potassium currents (I_KNa_) to 3 μM quinidine blockade in WT (**I**) and Na_V_1.6-KO (**J**) neurons. I_KNa_ before application of quinidine was obtained from the subtraction of I_Control_ and I_Li_. Maintained I_KNa_ after application of 3 μM quinidine was obtained from the subtraction of I_Quid_ and I_Li+Quid_. (**K**) Summarized amplitudes of I_KNa_ before and after application of 3 μM quinidine in the bath solution in WT (black, n = 12) and Na_V_1.6-KO (red, n = 10) primary cortical neurons. \*\*\*\**p* < 0.0001, Two-way repeated measures ANOVA followed by Bonferroni’s post hoc test.

Slack and Slick are both K_Na_ channels (with 74% sequence identity) and adopt similar structures^29^, and Slick is also blocked by quinidine^30^. We next assessed whether Na_V_1.6 sensitizes Slick to quinidine blockade and observed that, similar to Slack, co-expression of Slick and Na_V_1.6 in HEK293 cells resulted in a sensitization of Slick to quinidine blockade (7-fold) (Fig. S2A,B). These results support that Na_V_1.6 regulates both Slack and Slick and that Na_V_1.6 can sensitize K_Na_ channels to quinidine blockade *in vitro*.

We also asked whether Na_V_1.6 sensitizes native K_Na_ channels to quinidine blockade in neurons. We performed whole-cell patch-clamp recordings in primary cortical neurons from postnatal (P0-P1) homozygous Na_V_1.6 knockout C3HeB/FeJ mice and the wild-type littermate controls. As Lithium is a much weaker activator of K_Na_ channels than sodium^29^, K_Na_ currents (I_KNa_) were isolated by replacing sodium ions with equivalent lithium ions in the bath solution (Fig. 1G,H) as previously described^22^. The amplitudes of I_KNa_ remained unaffected by the homozygous knockout of Na_V_1.6 (Fig. S3A). Notably, 3 μM quinidine significantly inhibited native I_KNa_ (44%) in wild-type neurons (Fig. 1I,K) and this inhibitory effect was not limited to targeting the larger I_KNa_ (Fig. S3B). Conversely, the same concentration of quinidine had no significant effect on I_KNa_ in Na_V_1.6-knockout (Na_V_1.6-KO) neurons (Fig. 1J,K). These results support that Na_V_1.6 is required for the observed high sensitivity of native K_Na_ channels to quinidine blockade.

### Transient sodium influx through Na_V_1.6 enhances Na_V_1.6-mediated sensitization of Slack to quinidine blockade

We next investigated the biomolecular mechanism underlying Na_V_1.6-mediated sensitization of Slack to quinidine blockade. Considering that Slack currents are activated by sodium influx^22,29^, we initially assessed the effects of Na_V_1.6-mediated sodium influx on sensitizing Slack to quinidine blockade. We used 100 nM tetrodotoxin (TTX) to block Na_V_1.6-mediated sodium influx (Fig. S4A,B)^31^. In HEK293 cells expressing Slack alone, 100 nM TTX did not affect Slack currents; nor did it affect Slack’s sensitivity to quinidine blockade (IC_50_ = 83.27 μM) (Fig. 2A,B and Fig. S4C,D). In contrast, upon co-expression of Slack and Na_V_1.6 in HEK293 cells, bath-application of 100nM TTX significantly reduced the effects of Na_V_1.6 in sensitizing Slack to quinidine blockade (IC_50_ = 25.04 μM) (Fig. 2A,B). These findings support that sodium influx through Na_V_1.6 contributes to Na_V_1.6-mediated sensitization of Slack to quinidine blockade.

**Figure 2.**
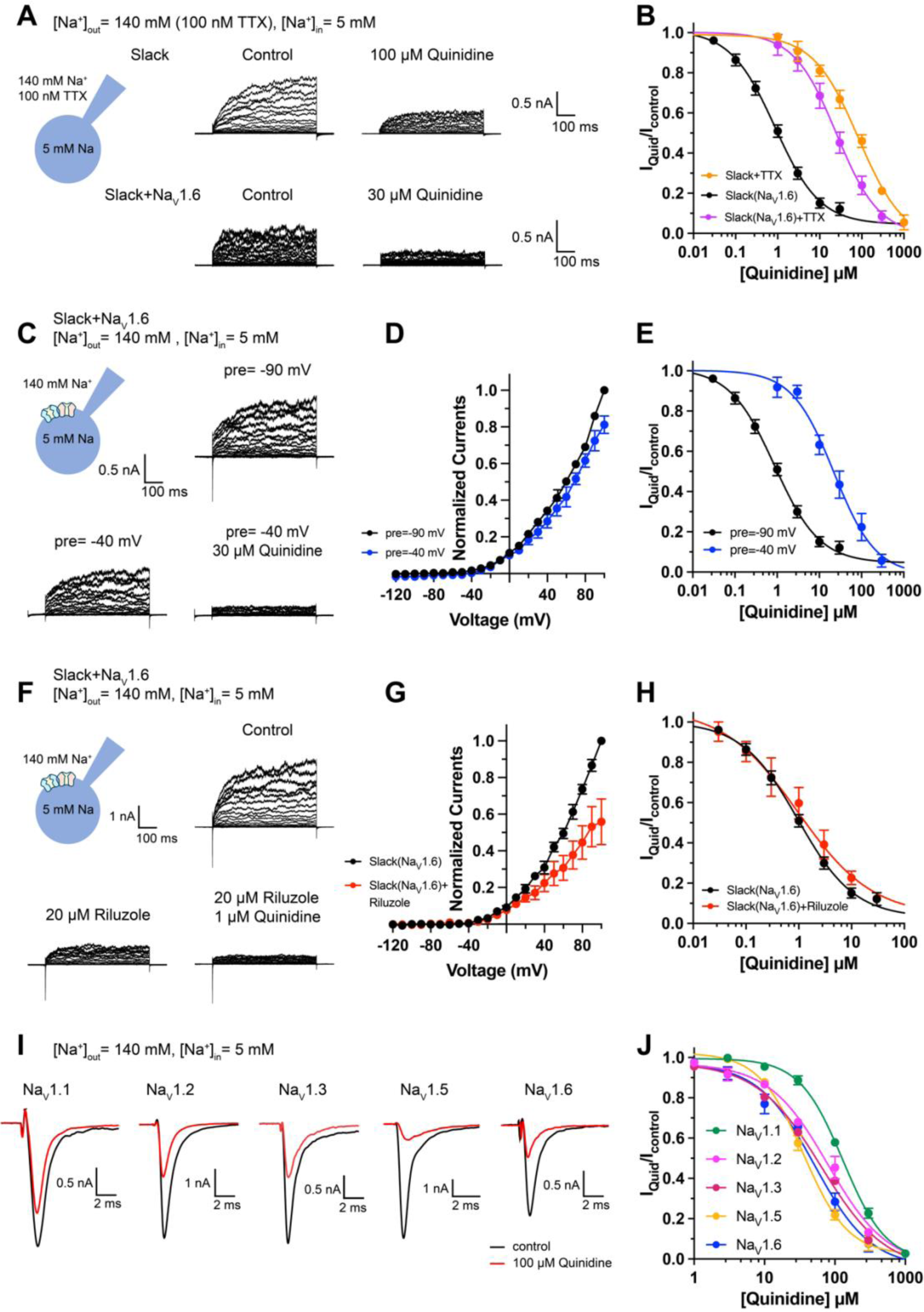
Blocking transient sodium influx through Na_V_1.6 reduces Na_V_1.6-mediated sensitization of Slack to quinidine blockade. (**A**) Example current traces from HEK293 cells expressing Slack alone (top) and co-expressing Slack with Na_V_1.6 (bottom), with 100nM TTX in the bath solution. The left traces show the family of control currents; the right traces show Slack currents remaining after application of quinidine. The presented concentrations of quinidine were chosen to be near the IC_50_ values. (**B**) The concentration-response curves for blocking of Slack by quinidine at +100 mV upon expression of Slack alone (n = 3) and co-expression of Slack with Na_V_1.6 (n = 7), with 100 nM TTX in the bath solution. (**C**) Top, example current traces recorded from a HEK293 cell co-expressing Slack with Na_V_1.6 and evoked from a 100-ms prepulse (pre) of -90 mV, with the same voltage protocol as in Fig. 1D. Bottom, example current traces recorded from the same cell but evoked from a 100-ms prepulse of -40 mV, before and after application of quinidine. (**D**) I-V curves of Slack upon co-expression with Na_V_1.6. The currents were evoked from a prepulse of -90 mV (black) or -40 mV (blue). (**E**) The concentration-response curves for blocking of Slack by quinidine at +100 mV with a prepulse of -90 mV (black, n = 19) or -40 mV (blue, n = 5). (**F**) Top, example current traces recorded from a HEK293 cell co-expressing Slack and Na_V_1.6 without riluzole in the bath solution. Bottom, example current traces recorded from the same cell with 20 µM riluzole in the bath solution, before and after application of quinidine. (**G**) I-V curves of Slack upon co-expression with Na_V_1.6 before (black) and after (red) application of 20 µM riluzole into bath solution.The concentration-response curves for blocking of Slack by quinidine upon co-expression of Slack with Na_V_1.6, without (n = 19) or with (n = 6) 20 µM riluzole in the bath solution. (**I-J**) The sensitivity of Na_V_ channel subtypes to quinidine blockade upon expression of Na_V_ alone in HEK293 cells. Example current traces (**I**) were evoked by a 50-ms step depolarization to 0 mV from a holding potential of -90 mV. The Concentration-response curves for blocking of Na_V_ channel subtypes by quinidine (**J**) were shown on the right panel (n = 5 for Na_V_1.1, n = 3 for Na_V_1.2, n = 6 for Na_V_1.3, n = 6 for Na_V_1.5, and n = 4 for Na_V_1.6).

It is known that Na_V_1.6-mediated sodium influx involves a transient inward flux that reaches a peak before subsequently decaying to the baseline within a few milliseconds; this is termed a transient sodium current (I_NaT_)^32^. A small fraction of Na_V_1.6 currents are known to persist after the rapid decay of I_NaT,_ and these are termed persistent sodium currents (I_NaP_)^33^. We isolated I_NaT_ and I_NaP_ to explore their potential contributions in sensitizing Slack to quinidine blockade. We selectively inactivated I_NaT_ using a depolarized prepulse of -40 mV (Fig. S5A) and selectively blocked I_NaP_ by bath-application of 20 μM riluzole, which is a relatively specific I_NaP_ blocker that is known to stabilize inactivated-state Na_V_ channels and delay recovery from inactivation^34,35^. Our findings ultimately confirmed that the 20 μM riluzole selectively blocked I_NaP_ compared to I_NaT_ in HEK293 cells co-expressing Slack and Na_V_1.6 (Fig. S5B) and that 20 μM riluzole had no effect on Slack currents when expressed alone (Fig. S5C,D).

Consistent with previous investigations^21,22^, inactivating I_NaT_ reduced whole-cell Slack currents by 20%, and blocking I_NaP_ reduced Slack currents by 40% at +100 mV (Fig. 2C,D,F,G), supporting that Na_V_1.6-mediated sodium influx activates Slack. Interestingly, inactivating I_NaT_ resulted in a > 20-fold decrease in Slack’s sensitization to quinidine blockade (IC_50_ = 22.26 μM) (Fig. 2C,E). In contrast, blocking I_NaP_ had no effect on Slack’s sensitization to quinidine blockade (IC_50_ = 1.60 μM) (Fig. 2F,H). These findings indicate that Na_V_1.6 sensitizes Slack to quinidine blockade via I_NaT_ but not I_NaP_.

Given that Slack current amplitudes are sensitive to sodium influx, and considering that quinidine is a sodium channel blocker, we examined whether Na_V_1.6 has higher sensitivity to quinidine blockade than other Na_V_ channel subtypes, which could plausibly explain the observed increased strength of sensitization. We used whole-cell patch-clamping to assess the sensitivity of Na_V_1.1, Na_V_1.2, Na_V_1.3, Na_V_1.5, and Na_V_1.6 to quinidine blockade. These sodium channels exhibited similar levels of quinidine sensitivity (IC_50_ values in the range of 35.61-129.84 μM) (Fig. 2I,J and Supplementary Table 2), all of which were at least 40-fold lower than the Na_V_1.6-mediated sensitization of Slack to quinidine blockade (Fig. 1F). Additionally, co-expressing Slack with Na_V_1.1, Na_V_1.2, Na_V_1.3, Na_V_1.5, or Na_V_1.6 in HEK293 cells did not change the sensitivity of these Na_V_ channel subtypes to quinidine blockade (Fig. S6 and Supplementary Table 2). Thus, differential quinidine affinity for specific Na_V_ channel subtypes cannot explain the large observed Na_V_1.6-mediated sensitization of Slack to quinidine blockade. Moreover, it is clear that Na_V_1.6-mediated sensitization of Slack to quinidine blockade is directly mediated by I_NaT_, rather than through some secondary effects related to Na_V_1.6’s higher sensitivity to quinidine blockade.

### Slack physically interacts with Na_V_1.6

We found that the specific voltage-gated sodium channel blocker TTX did not completely abolish the effects of Na_V_1.6 on sensitizing Slack to quinidine blockade (Fig. 2B), so it appears that a sodium-influx-independent mechanism is involved in the observed Na_V_1.6-mediated sensitization of Slack to quinidine blockade. We therefore investigated a potential physical interaction between Slack and Na_V_1.6. We initially assessed the cellular distribution of Na_V_1.2, Na_V_1.6, and Slack in the hippocampus and the neocortex of mouse. Consistent with previous studies^36,37^, Na_V_1.2 and Na_V_1.6 were localized to the axonal initial segment (AIS) of neurons, evident as the co-localization of Na_V_ and AnkG, a sodium channel-associated protein known to accumulate at the AIS (Fig. 3A). Slack channels were also localized to the AIS of these neurons (Fig. 3A), indicating that Slack channels are located in close proximity to Na_V_1.6 channels, and supporting their possible interaction. Moreover, Na_V_1.6 was co-immunoprecipitated with Slack in homogenates from mouse cortical and hippocampal tissues and from HEK293T cells co-transfected with Slack and Na_V_1.6 (Fig. 3B,C), supporting that Slack and Na_V_1.6 form protein complexes in mouse brains.

**Figure 3.**
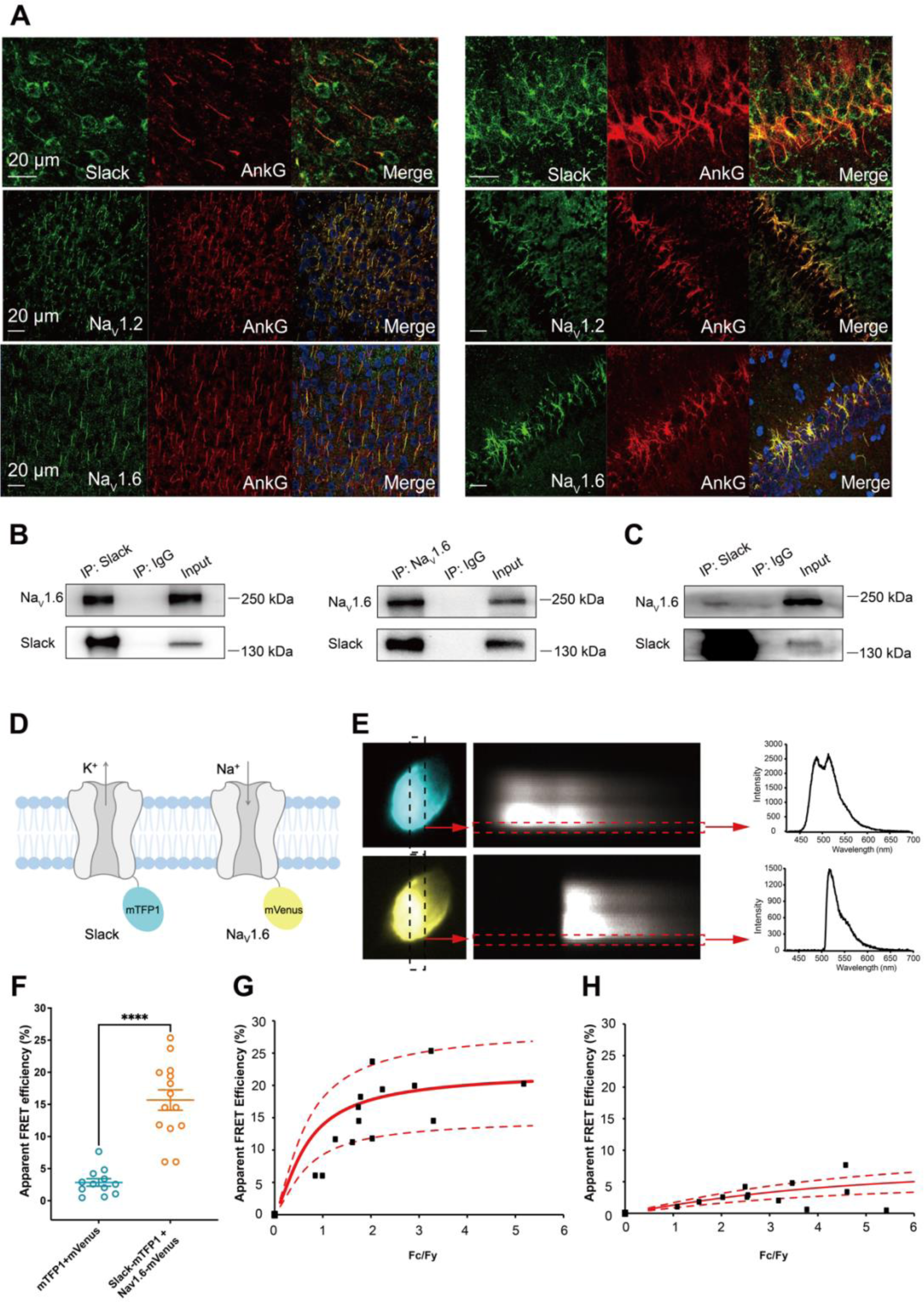
Slack physically interacts with Na_V_1.6. (**A**) Immunofluorescence of Slack, Na_V_1.2, Na_V_1.6 (green), and AnkG (red) in neocortex layer 5 (left) and hippocampal CA1 pyramidal cell layer (right). Confocal microscopy images were obtained from Coronal brain slices of C57BL/6 mice. The panels from top to bottom show the double staining of Slack with AnkG, Na_V_1.2 with AnkG, and Na_V_1.6 with AnkG, respectively. DAPI (blue) was used for nuclear counterstaining. (**B**) Coimmunoprecipitation (Co-IP) of Slack and Na_V_1.6 in cell lysates from HEK293T cells co-transfected with Slack and Na_V_1.6. (**C**) Co-IP of Slack and Na_V_1.6 in mouse brain tissue lysates. Input volume corresponds to 10% of the total lysates for Co-IP. (**D**) A schematic diagram showing the fluorescence-labeled Slack and Na_V_1.6. mTFP1 and mVenus were fused to the C-terminal region of Slack (Slack-mTFP1) and Na_V_1.6 (Na_V_1.6-mVenus), respectively. (**E**) FRET imaging of Slack-mTFP1 and Na_V_1.6-mVenus co-expressed in HEK293 cells. The emission spectra measured from the edge of cell (dotted arrows in red) are used for FRET efficiency calculation. (**F**) The apparent FRET efficiency measured from cells co-expressing the fluorophore-tagged ion channels or co-expressing the fluorophores. **** *p* < 0.0001, Mann-Whitney test. (**G-H**) The FRET efficiency measured from cells co-expressing the fluorophore-tagged ion channels (**G**), or from cells co-expressing fluorophores (**H**). The efficiency value was plotted as a function of the fluorescence intensity ratio between mTFP1 and mVenus (Fc/Fy). Each symbol represents a single cell. The solid curve represents the FRET model that yields the best fit; dotted curves represent models with 5% higher or lower FRET efficiencies.

To assess the interaction between Slack and Na_V_1.6 inside living cells, we performed a FRET assay in transfected HEK293T cells^38^. Briefly, we genetically fused mTFP1 and mVenus to the C-terminal regions of Slack and Na_V_1.6, respectively (Fig. 3D). Upon imaging the emission spectra cells co-expressing Na_V_1.6-mVenus and Slack-mTFP1 (measured at the plasma membrane region) (Fig. 3E), we detected positive FRET signals, indicating a Slack-Na_V_1.6 interaction (Fig. 3F,H). The plasma membrane regions from HEK293T cells co-transfected with Na_V_1.6 and Slack showed FRET efficiency values much larger than a negative control (in which standalone mVenus and mTFP1 proteins were co-expressed) (Fig. 3G,H), indicating that Slack channels reside in close spatial proximity (less than 10 nm) to Na_V_1.6 channels in living cells.

We next characterized the consequences of the Slack-Na_V_1.6 interaction in HEK293 cells using whole-cell recordings. Slack increased the rate of recovery from fast inactivation of Na_V_1.6 (Fig. S7E), with no significant effects on the steady-state activation, steady-state fast inactivation, or ramp currents (Fig. S7C,D,F and Supplementary Table 3). Additionally, we found that Na_V_1.6 had no significant effects on the activation rate or the current-voltage (I-V) relationship of Slack currents (Fig. S7A,B). These results indicate that the physical interaction between Slack and Na_V_1.6 produces functional consequences. Taken together, these findings support functional and physical coupling of Slack and Na_V_1.6.

### Na_V_1.6’s N- and C-termini bind to Slack’s C-terminus and sensitize Slack to quinidine blockade

To explore whether the physical interaction between Slack and Na_V_1.6 is required for sodium-influx-mediated sensitization of Slack to quinidine blockade, we performed inside-out patch-clamp recordings on HEK293 cells transfected with Slack alone or co-transfected with Slack and Na_V_1.6. Note that in these experiments the intracellular sodium concentration ([Na^+^]_in_) was raised to 140 mM (a concentration at which most Slack channels can be activated^20^), seeking to mimic the increased intracellular sodium concentration upon sodium influx. When expressing Slack alone, increasing the sodium concentration did not sensitize Slack to quinidine blockade (IC_50_ = 120.42 μM) (Fig. 4a). However, upon co-expression of Na_V_1.6 and Slack, Na_V_1.6 significantly sensitized Slack to quinidine blockade (IC_50_ = 2.91 μM) (Fig. 4A). These results support that physical interaction between Slack and Na_V_1.6 is a prerequisite for sodium-influx-mediated sensitization of Slack to quinidine blockade.

**Figure 4.**
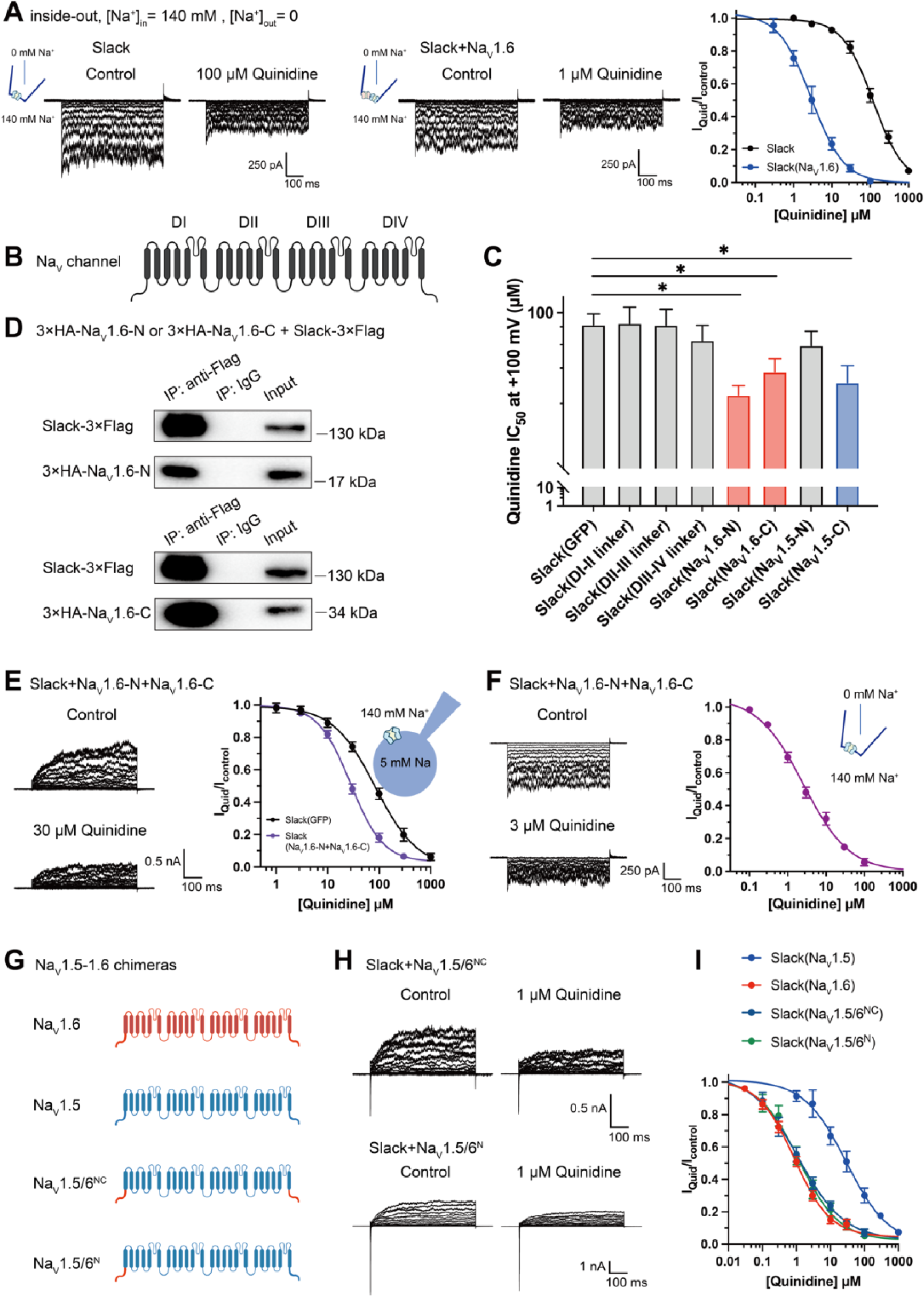
Na_V_1.6’s N- and C-termini interacting with Slack is a prerequisite for Na_V_1.6-mediated sensitization of Slack to quinidine blockade. (**A**) The sensitivity of Slack to quinidine blockade upon expression of Slack alone (n = 3) and co-expression of Slack with Na_V_1.6 (n = 3) from excised inside-out patches. The pipette solution contained (in mM) 130 KCl, 1 EDTA, 10 HEPES and 2 MgCl_2_ (pH 7.3); the bath solution contained (in mM) 140 NaCl, 1 EDTA, 10 HEPES and 2 MgCl_2_ (pH 7.4). The membrane voltage was held at 0 mV and stepped to voltages varying from −100 mV to 0 mV in 10 mV increments. Example current traces were shown on the left panel. The concentration-response curves for blocking of Slack by quinidine were shown on the right panel. (**B**) Domain architecture of the human Na_V_ channel pore-forming α subunit. (**C**) Calculated IC_50_ values at +100 mV of quinidine on Slack upon co-expression with indicated cytoplasmic fragments from Na_V_ channels. For Na_V_1.6, cytoplasmic fragments used include N-terminus (Na_V_1.6-N, residues 1-132), inter domain linkers (Domain I-II linker, residues 409-753; Domain II-III linker, residues 977-1199; Domain III-IV linker, residues 1461-1523), and C-terminus (Na_V_1.6-C, residues 1766-1980). For Na_V_1.5, cytoplasmic fragments used include N-terminus (Na_V_1.5-N, residues 1-131) and C-terminus (Na_V_1.5-C, residues 1772-2016). (**D**) Co-IP of Slack and terminal domains of Na_V_1.6 in cell lysates from HEK293T cells co-expressing 3×Flag-tagged Slack (Slack-3×Flag) and 3×HA-tagged termini of Na_V_1.6 (3×HA-Na_V_1.6-N or 3×HA-Na_V_1.6-C). The 3×Flag tag was fused to the C-terminal region of Slack and the 3×HA tag was fused to the N-terminal region of Na_V_1.6’s fragments. (**E**) The sensitivity of Slack to quinidine blockade upon co-expression of Slack with GFP (n = 12) or N- and C-termini of Na_V_1.6 (n = 11), from whole-cell recordings. (**F**) The sensitivity of Slack to quinidine blockade upon co-expression of Slack with N- and C-termini of Na_V_1.6, from excised inside-out recordings (n = 10, using the same protocols as in Fig. 4A). Example current traces before and after application of quinidine were shown on the left panel. The concentration-response curves were shown on the right panel. (**G**) A schematic diagram of the Na_V_1.5-1.6 chimeric channels (Na_V_1.5/6^NC^ and Na_V_1.5/6^N^) used in this study. (**H**) Example current traces recorded from HEK293 cells co-expressing Slack and Na_V_1.5-1.6 chimeras before and after application of the indicated concentration of quinidine. (**I**) The concentration-response curves for blocking of Slack by quinidine upon co-expression of Slack with Na_V_1.5 (n = 9), Na_V_1.6 (n = 19), Na_V_1.5/6^NC^ (n = 9), or Na_V_1.5/6^N^ (n = 9).

To investigate which interacting domains mediate Na_V_1.6’s sensitization of Slack to quinidine blockade, we focused on Na_V_1.6’s cytoplasmic fragments, including its N-terminus, inter domain linkers, and C-terminus (Fig. 4B). Whole-cell recordings from HEK293 cells co-expressing Slack with these Na_V_1.6 fragments revealed that the N-terminus and C-terminus of Na_V_1.6 significantly enhanced the sensitivity of Slack to quinidine blockade (IC_50_ = 31.59 μM for Slack upon co-expression with Na_V_1.6’s N-terminus and IC_50_ = 43.70 μM for Slack upon co-expression with Na_V_1.6’s C-terminus); note that the inter domain linkers had no effect (Fig. 4C).

Subsequent co-immunoprecipitation and glutathione S-transferase (GST) pull down assays of HEK293T cell lysates experimentally confirmed that Na_V_1.6’s N-and C-termini each interact with Slack (Fig. 4D and Fig. S8). Additionally, whole-cell recordings using an [Na^+^]_in_ of 5 mM again showed that Na_V_1.6’s N-terminus and Na_V_1.6’s C-terminus sensitize Slack to quinidine blockade (IC_50_ = 27.87 μM) (Fig. 4E). And inside-out recordings using an [Na^+^]_in_ of 140 mM showed that co-expression of Slack, Na_V_1.6’s N-terminus, and Na_V_1.6’s C-terminus resulted in obvious sensitization of Slack to quinidine blockade, with an IC_50_ of 2.57 μM (Fig. 4F)—thus fully mimicking the aforementioned effects of full-length Na_V_1.6 (IC_50_ = 2.91 μM) (Fig. 4A). These findings support that the binding of Na_V_1.6’s N- and C-termini to Slack is required for Na_V_1.6’s sensitization of Slack to quinidine blockade.

Recalling that Na_V_1.5 had the least pronounced effect in sensitizing Slack to quinidine blockade among all examined Na_V_ channels (IC_50_ = 29.46 μM) (Fig. 1F and Supplementary Table 1), we initially compared the sodium currents of Na_V_ channel subtypes co-expressed with Slack. The results revealed no significant differences in activation time constants (Fig. S9A). It’s worth noting that Na_V_1.5 exhibited significantly larger current amplitudes than Na_V_1.6 (Fig. S9B). Subsequently, we constructed Na_V_1.5-1.6 chimeras to test the roles of Na_V_1.6’s N- and C-termini in sensitizing Slack to quinidine blockade. The replacement of both the N-terminus [residues 1-131] and C-terminus [residues 1772-2016] of Na_V_1.5 with Na_V_1.6’s N-terminus [residues 1-132] and C-terminus [residues 1766-1980] (namely Na_V_1.5/6^NC^) fully mimicked effects of Na_V_1.6 in sensitizing Slack to quinidine blockade (IC_50_ = 1.13 μM) (Fig. 4G-I). We also found that replacement of Na_V_1.5’s N-terminus [residues 1-131] with Na_V_1.6’s N-terminus [residues 1-132] (namely Na_V_1.5/6^N^) fully mimicked the effects of Na_V_1.6 (IC_50_ = 1.18 μM) (Fig. 4G-I).

Notably, despite both Na_V_1.5 and Na_V_1.5/6^N^ exhibited much larger current amplitudes compared to Na_V_1.6 (Fig. S9B), only Na_V_1.5/6^N^ replicated the effect of Na_V_1.6 in sensitizing Slack to quinidine blockade (Fig. 4H-I). These results suggest that the observed differences between Na_V_1.5 and Na_V_1.6 in sensitizing Slack are unlikely to be attributed to Na_V_1.6’s smaller sodium current amplitudes but involve Na_V_1.6’s N-terminus. Consistently, Na_V_1.5’s C-terminus sensitized Slack to quinidine blockade (IC_50_ = 37.59 μM), whereas Na_V_1.5’s N-terminus had no effect on Slack sensitization (Fig. 4C). These findings support that Na_V_1.6’s N-terminus is essential for sensitizing Slack to quinidine blockade.

Having demonstrated that Na_V_1.6 sensitizes Slack via Na_V_1.6’s cytoplasmic N- and C-termini, we investigated which domains of Slack interact with Na_V_1.6 and focused on Slack’s cytoplasmic fragments, including Slack’s N-terminus and C-terminus (Fig. 5A). In HEK293 cells co-expressing Na_V_ and Slack, Na_V_-mediated sensitization of Slack to quinidine blockade was significantly attenuated upon the additional expression of Slack’s C-terminus, but not of Slack’s N-terminus (Fig. 5B,C), suggesting that Slack’s C-terminus can disrupt the Slack-Na_V_1.6 interaction by competing with Slack for binding to Na_V_1.6. Consistently, Slack’s C-terminus co-immunoprecipitated with Na_V_1.6’s N- and C-termini in HEK293T cell lysates (Fig. 5D). Together, these results support that Slack’s C-terminus physically interacts with Na_V_1.6 and that this interaction is required for Na_V_1.6’s sensitization of Slack to quinidine blockade.

**Figure 5.**
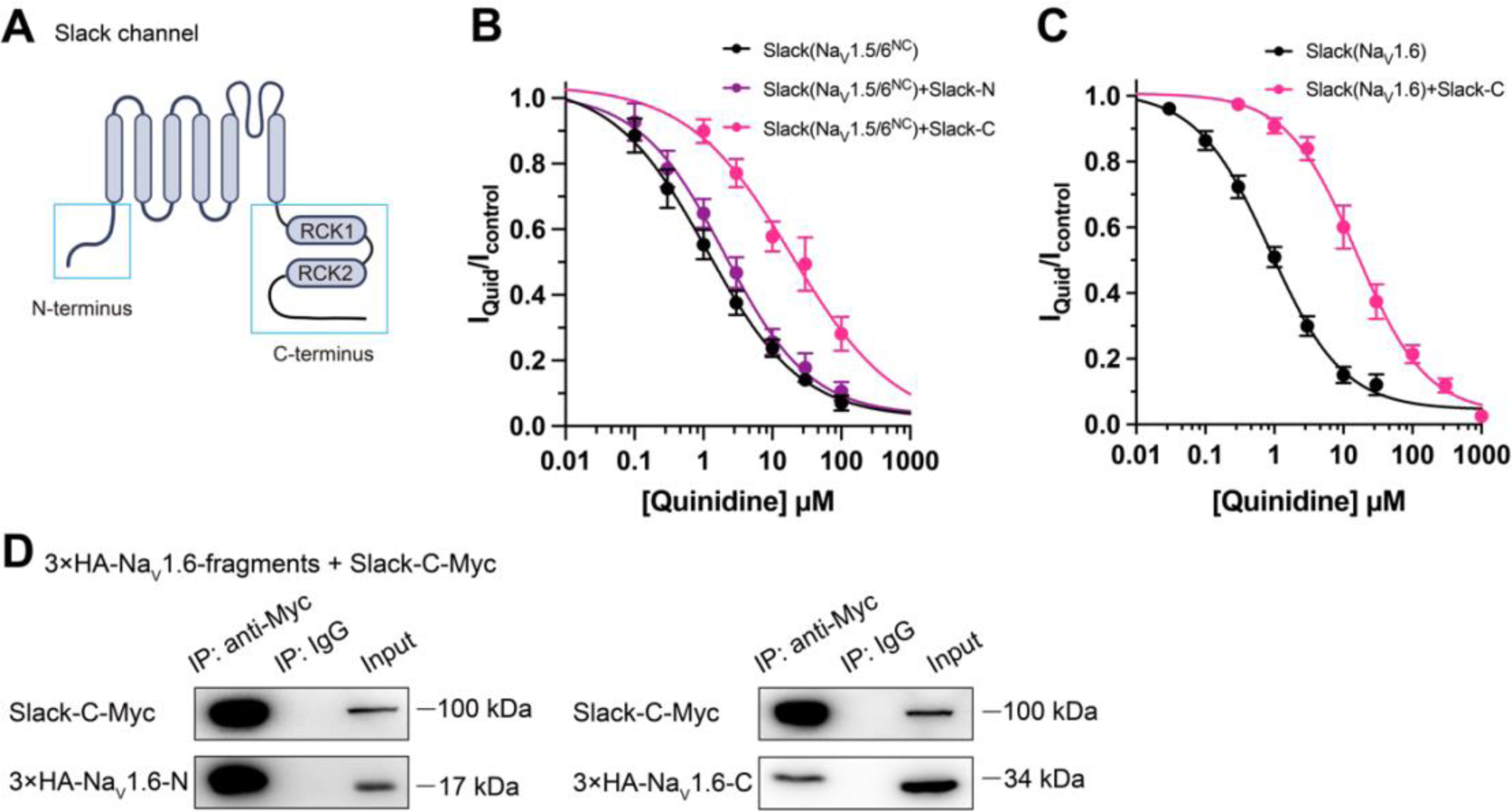
Slack’s C-terminus is required for Na_V_1.6-mediated sensitization of Slack to quinidine blockade. (**A**) Domain architecture of the human Slack channel subunit. Slack’s N-terminus (Slack-N, residues 1-116) and C-terminus (Slack-C, residues 345-1235) were shown in the blue boxes. (**B**) The concentration-response curves for blocking of Slack by quinidine upon additional expression of Slack’s N- or C-terminus in HEK293T cells co-expressing Slack and Na_V_1.5/6^NC^. (**C**) The concentration-response curves for blocking Slack by quinidine upon additional expression of Slack’s C-terminus in HEK293 cells co-expressing Slack and Na_V_1.6. (**D**) Co-IP of Myc-tagged Slack’s C-terminus (Slack-C-Myc) with 3×HA-tagged Na_V_1.6’s termini (3×HA-Na_V_1.6-N or 3×HA-Na_V_1.6-C) in HEK293T cell lysates. The 3×HA tag was fused to the N-terminal region of Na_V_1.6’s fragments, and the Myc tag was fused to the C-terminal region of Slack’s fragment.

### Na_V_1.6 binds to and sensitizes epilepsy-related Slack mutant variants to quinidine blockade

Over 50 mutations in KCNT1 (Slack) have been identified and related to seizure disorders^11^. Having established that Na_V_1.6 can sensitize wild-type Slack to quinidine blockade, we next investigated whether Na_V_1.6 also sensitizes epilepsy-related Slack mutant variants to quinidine blockade. We chose 3 Slack pathogenic mutant variants (K629N, R950Q, and K985N) initially detected in patients with KCNT1-related epilepsy^15,39,40^. Considering that these 3 mutations are located in Slack’s C-terminus, and recalling that Slack’s C-terminus interacts with Na_V_1.6 (Fig. 5D), we first used co-immunoprecipitation assays and successfully confirmed that each of these Slack mutant variants interacts with Na_V_1.6 in HEK293T cell lysates (Fig. 6A).

**Figure 6.**
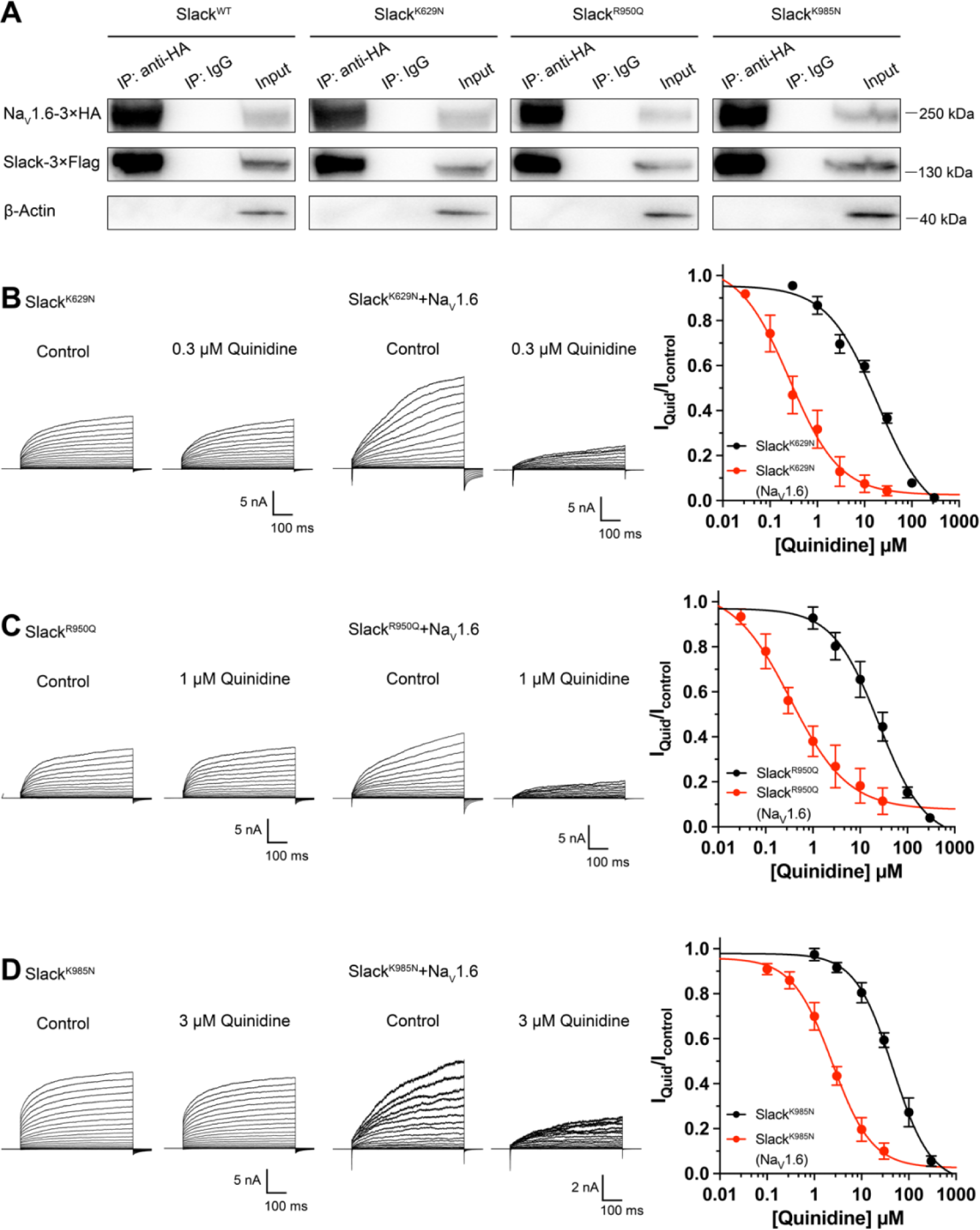
Na_V_1.6 sensitizes epilepsy-related Slack mutant variants to quinidine blockade. (**A**) Co-IP of 3×Flag-tagged Slack or its mutations (Slack-3×Flag) with 3×HA-tagged Na_V_1.6 (Na_V_1.6-3×HA) in HEK293T cell lysates. The tags were all fused to the C-terminal region of wild-type or mutant ion channels. (**B-D**) The sensitivity of Slack mutant variants (Slack^K629N^ [**B**], Slack^R950Q^ [**C**], and Slack^K985N^ [**D**]) to quinidine blockade upon expression of Slack mutant variants alone and co-expression of Slack mutant variants with Na_V_1.6. Left, example current traces recorded from HEK293 cells expressing Slack mutant variants alone and co-expressing Slack mutant variants with Na_V_1.6, before and after application of the indicated concentrations of quinidine. Right, the concentration-response curves for blocking of Slack mutant variants by quinidine upon expression of Slack mutant variants alone (n = 8 for Slack^K629N^, n = 7 for Slack^R950Q^, and n = 5 for Slack^K985N^) and co-expression of Slack mutant variants with Na_V_1.6 (n = 8 for Slack^K629N^ upon co-expression with Na_V_1.6, n = 5 for Slack^R950Q^ upon co-expression with Na_V_1.6, and n = 7 for Slack^K985N^ upon co-expression with Na_V_1.6). Please refer to Supplementary Table 4 for IC_50_ values.

Subsequently, whole-cell recordings revealed that the examined Slack mutant variants exhibited no discernible impact on the amplitudes of Na_V_1.6 currents (Fig. S10A), while pharmacologically blocking Na_V_1.6 currents using bath-application of 100 nM TTX significantly reduced the amplitudes of Slack mutant variant (Slack^R950Q^) currents (Fig. S10B), suggesting a Na^+^-mediated functional coupling between Slack mutant variant and Na_V_1.6 currents. Notably, Na_V_1.6 significantly sensitized all of the examined Slack mutant variants to quinidine blockade, with IC_50_ values ranging from 0.26 to 2.41 μM (Fig. 6B-D and Supplementary Table 4). These results support that Na_V_1.6 interacts with examined Slack mutant variants and sensitizes them to quinidine blockade. It is plausible that the Slack-Na_V_1.6 interaction contributes to the therapeutical role of quinidine in the treatment of KCNT1-related epilepsy.

### Viral expression of Slack’s C-terminus prevents Slack^G269S^-induced seizures

Having established that blocking Na_V_1.6-mediated sodium influx significantly reduced Slack current amplitudes (Fig. 2D,G and Fig. S10B), we found that the heterozygous knockout of Na_V_1.6 significantly reduced the afterhyperpolarization amplitude in murine hippocampal neurons (Fig. S11), together indicating that Na_V_1.6 activates native Slack through providing Na^+^. We therefore assumed that disruption of the Slack-Na_V_1.6 interaction should reduce the amount of Na^+^ in the close vicinity of epilepsy-related Slack mutant variants and thereby counter the increased current amplitudes of these Slack mutant variants. Pursuing this, we disrupted the Slack-Na_V_ interaction by overexpressing Slack’s C-terminus (to compete with Slack) and measured whole-cell current densities. In HEK293 cells co-expressing epilepsy-related Slack mutant variants (G288S, R398Q)^41,42^ and Na_V_1.5/6^NC^, expression of Slack’s C-terminus significantly reduced whole-cell current densities of Slack^G269S^ and Slack^R398Q^ (Fig. 7A,B), supporting that disrupting the Slack-Na_V_1.6 interaction can indeed reduce current amplitudes of Slack mutant variants, which may protect against seizures induced by Slack mutant variants.

**Figure 7.**
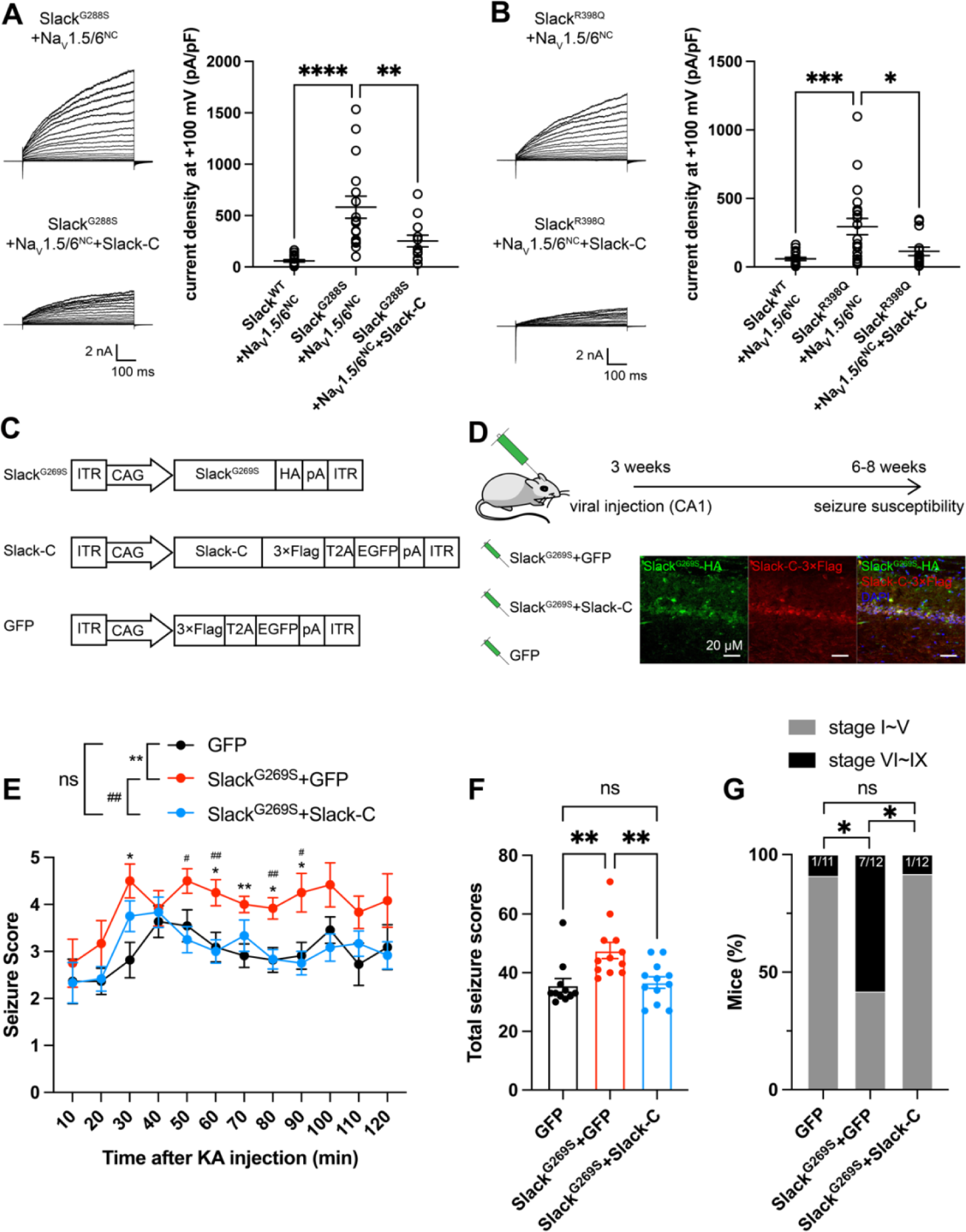
Viral expression of Slack’s C-terminus prevents Slack^G269S^-induced seizures. (**A-B**) The current densities of Slack mutant variants (Slack^G288S^ [**A**] and Slack^R398Q^ [**B**]) upon co-expression with Na_V_1.5/6^NC^ in HEK293T cells were reduced by additional expression of Slack’s C-terminus. Left, example current traces from HEK293T cells co-expressing Slack mutant variants and Na_V_1.5/6^NC^ or co-expressing Slack mutant variants, Na_V_1.5/6^NC^, and Slack’s C-terminus. Right, summarized current densities at +100 mV. * *p* < 0.05, ** *p* < 0.01, *** *p* < 0.001; one-way ANOVA followed by Bonferroni’s post hoc test. (**C**) Architecture for expression cassettes of AAVs. (**D**) Top, study design and timeline for the stereotactic injection model. Bottom, Immunofluorescence of HA-tagged Slack^G269S^ (green), 3×Flag-tagged Slack’s C-terminus (red), and DAPI (blue) in hippocampal CA1 pyramidal cell layer at 5 weeks after viral injection of Slack^G269S^ with Slack’s C-terminus into CA1 of mice. (**E**) Time-course of KA-induced seizure stage changes at 10-min intervals based on a modified Racine, Pinal, and Rovner scale (please refer to Methods for further details). The number of mice used: “GFP” control group (n = 11), “Slack^G269S^+GFP” group (n = 12), “Slack^G269S^+Slack-C” group (n = 12). “GFP” vs. “Slack^G269S^+GFP”: F_(1,21)_ = 10.48, *p* = 0.0040, * *p* < 0.05, ** *p* < 0.01; “Slack^G269S^+GFP” vs. “Slack^G269S^+Slack-C”: F_(1,22)_ = 10.30, *p* = 0.0040, ^#^ *p* < 0.05, ^##^ *p* < 0.01. “GFP” vs. “Slack^G269S^+Slack-C”: F_(1,21)_ = 0.09574, *p* = 0.7600. Repeated two-way ANOVA followed by Bonferroni’s post hoc test. (**F**) Total seizure score per mouse over the 2 h after KA injection of these three groups. * *p* < 0.05, ** *p* < 0.01; one-way ANOVA followed by Bonferroni’s post hoc test. (**G**) The percentage of mice with stage VI∼IX seizures over the 2 h after KA injection in each group. * *p* < 0.05; Fisher’s exact test.

We next induced an *in vivo* epilepsy model by introducing a Slack G269S variant into C57BL/6N mice using adeno-associated virus (AAV) injection to mimic the human Slack mutation G288S^41^. Specifically, we delivered stereotactic injections of AAV9 containing expression cassettes for Slack^G269S^ (or GFP negative controls) into the hippocampal CA1 region of 3-week-old C57BL/6N mice (Fig. 7C,D). At 3-5 week intervals after AAV injection, we quantified the seizure susceptibility of mice upon the induction of a classic kainic acid (KA) model of temporal lobe epilepsy^43,44^. In this model, seizures with stage IV or higher (as defined by a modified Racine, Pinal, and Rovner scale^45^) are induced in rodents by intraperitoneal administration of 28 mg/kg KA.

We assessed a time course of KA-induced seizure stages at 10-min intervals and found that viral expression of Slack^G269S^ resulted in faster seizure progression in mice compared to control GFP expression (Fig. 7E). We calculated the total seizure score per mouse to assess seizure severity^46^. Slack^G269S^-expressing mice showed significantly higher seizure severity than GFP-expressing mice (Fig. 7F). The percentage of mice with stage VI∼IX seizures also increased, from 9.1% in GFP-expressing control mice to 58.3% in the Slack^G269S^-expressing mice (Fig. 7G). These results support that viral expression of Slack^G269S^ significantly increases seizure susceptibility in mice.

To evaluate the potential therapeutic effects of disrupting the Slack-Na_V_1.6 interaction, we delivered two AAV9s (one for Slack^G269S^ and one for Slack’s C-terminus [residues 326-1238]) into the CA1 region of mice (Fig. 7D). Viral expression of Slack’s C-terminus in Slack^G269S^-expressing mice significantly decreased seizure progression, seizure severity, and the percentage of mice experiencing stage VI∼IX seizures (Fig. 7E-G). These results support that viral expression of Slack’s C-terminus can prevent Slack^G269S^-induced seizures in mice, thus showcasing that using Slack’s C-terminus to disrupt the Slack-Na_V_1.6 interaction is a promising therapeutic intervention to treat KCNT1-related epilepsy.

## DISCUSSION

We here found that Na_V_1.6’s N- and C-termini bind to Slack’s C-terminus and sensitize Slack to quinidine blockade via Na_V_1.6-mediated transient sodium currents. These results suggest that the pharmacological blocking effects of a channel blocker are not exclusively mediated by the channel *per.se*., but modulated by channel’s interacting proteins. Moreover, we show that viral expression of Slack’s C-terminus can rescue the increased seizure susceptibility and confer protection against Slack^G269S^-induced seizures in mice.

At resting membrane potential, the intracellular sodium concentration ([Na^+^]_in_) in neurons (equal to ∼10 mM) is too low to effectively activate Slack (K_d_ of 66 mM)^1,20^. Slack is functionally coupled to sodium influx, which is known to be mediated by ion channels and receptors, including Na_V_, AMPARs, and NMDARs^21–23,47^. Such Na^+^ sources can provide both the membrane depolarization and the Na^+^ entry known to be required for Slack activation, enabling Slack to contribute both to action potential repolarization during neuronal high-frequency firing^7,48^ and to regulating excitatory postsynaptic potential (EPSP) at post-synaptic neurons^23^. Our results support that Slack and Na_V_1.6 form a channel complex, while also implying that Na_V_1.6-mediated sodium influx increases the Na^+^ concentration in the close vicinity of Slack to activate Slack. As a low threshold Na_V_ channel subtype, Na_V_1.6 has been reported to dominate the initiation and propagation of action potentials in axon initial segments (AIS) in excitatory neurons^36,49–51^. The activation of Slack by Na_V_1.6 at AIS has multiple impacts, including ensuring the timing of the fast-activated component of Slack currents, regulating the action potential amplitude, and apparently contributing to intrinsic neuronal excitability^29^. The use of HEK cells and cultured primary cortical neurons in this study may not fully capture the complexity of native Slack-Na_V_1.6 interaction. Future studies employing more intact systems, such as *in vivo* models, could provide a more comprehensive understanding of the physiological relevance of the native Slack-Na_V_1.6 interaction.

An interesting question arises from our observation that Slack’s sensitivity to quinidine blockade is enhanced by I_NaT_ but not I_NaP_: what can explain the distinct I_NaT_ and I_NaP_ contributions? We speculate that with physical modulation by Na_V_1.6, I_NaT_ may elicit a specific open conformation of Slack that brings its quinidine binding pocket into a high affinity state, which could lead to a substantial increase in Slack’s sensitivity to quinidine blockade. The possibility of this hypothetical open conformation is supported by previous reports of the presence of subconductance states detected in single-channel recordings of *Xenopus* oocytes expressing Slack channels^52^, which implies that there are multiple open conformations of Slack. Although only one open conformation of Slack has been observed in cryo-EM, the gap between maximum conformational open probability (∼1.0) and maximum functional open probability (∼0.7) implies a subclass within this class of open channels^53^. Additionally, it’s worth noting that blocking of I_NaP_ by application of riluzole has its limitation on selectivity. Riluzole has been reported to bind to Slack channels with low affinity^54^. Therefore, the dual binding of riluzole to both Slack and Na_V_1.6 may stabilize or inhibit the Slack-Na_V_1.6 interaction, consequently influencing the sensitization of Slack to quinidine blockade.

As previously mentioned, gain-of-function Slack mutant variants have been linked to a broad spectrum of epileptic disorders that are accompanied by intellectual disabilities and both psychomotor and developmental defects^18,55^. Given that many patients are refractory or non-responsive to conventional anticonvulsants^12,55,56^, and considering the limited success of quinidine in clinical treatment of KCNT1-related epilepsy, inhibitors targeting Slack are needed urgently^18^. Several small-molecule inhibitors against Slack have been reported, providing informative starting points for drug development efforts with KCNT1-related epilepsy^28,57,58^. However, our discovery of the Slack-Na_V_1.6 complex challenges the traditional view that Slack acts as an isolated target in KCNT1-related epilepsy^18^. Indeed, our study supports that co-expression of Slack and Na_V_1.6 in heterologous cell models should be performed when analyzing clinically relevant Slack mutations and when screening anti-epileptic drugs for use in treating KCNT1-related disorders.

Genotype-phenotype analysis has shown that ADSHE-related mutations are clustered in the regulator of conductance of K^+^ (RCK2) domain; while EIMFS-related mutations do not show a particular pattern of distribution^11^. All functionally tested Slack mutant variants show gain-of-function phenotypes, with increased Slack currents ^9,11,18^. Further, the epilepsy-related Slack mutant variants confer their gain-of-function phenotypes through two molecular mechanisms: increasing maximal channel open probability (P_max_) or increasing sodium sensitivity (K_d_) of Slack^59^. Both P_max_ and K_d_ are highly sensitive to [Na^+^]_in_; these are respectively analogous to the efficacy and potency of [Na^+^]_in_ on Slack currents^59^. Notably, several Slack mutant variants show gain-of-function phenotypes only at high [Na^+^]_in_ (*e.g.* 80 mM)^59^. These results indicate that the gain-of-function phenotype of epilepsy-related Slack mutant variants is aggravated by high [Na^+^]_in_. Our discovery of functional coupling between Na_V_1.6 and Slack presents a plausible basis for how Na_V_1.6-mediated sodium influx can increase [Na^+^]_in_ and thus apparently aggravate the gain-of-function phenotype of Slack mutant variants. Therefore, it makes sense that disruption of the Slack-Na_V_1.6 interaction by overexpressing Slack’s C-terminus reduces the current amplitudes of gain-of-function Slack mutant variants (Fig. 7A,B). Our successful demonstration that viral expression of Slack’s C-terminus prevents epilepsy-related Slack^G269S^-induced seizures in mice (Fig. 7E-G) warrants further translational evaluation for developing therapeutic interventions to treat KCNT1-related epilepsy.

## METHODS

### Animals

C57BL/6 mice were purchased from Charles River Laboratories. Na_V_1.6 knockout C3HeB/FeJ mice were generous gifts from Professor Yousheng Shu at Fudan University. All animals were housed on a 12-hour light/dark cycle with *ad libitum* access to food and water. All procedures related to animal care and treatment were approved by the Peking University Institutional Animal Care and Use Committee and met the guidelines of the National Institute of Health Guide for the Care and Use of Laboratory Animals. Each effort was made to minimize animal suffering and the number of animals used. The experiments were blind to viral treatment condition during behavioral testing.

### Antibodies and Reagents

Commercial antibodies used were: anti-AnkG (Santa Cruz), anti-Slack (NeuroMab), anti-Na_V_1.2 (Alomone), anti-Na_V_1.6 (Alomone), anti-HA (Abbkine), anti-Flag (Abbkine), anti-β-Actin (Biodragon), HRP Goat Anti-Mouse IgG LCS (Abbkine), HRP Mouse Anti-Rabbit IgG LCS (Abbkine), Alexa Fluor 488-AffinityPure Fab Fragment Donkey anti-rabbit IgG (Jackson), Alexa Fluor 594 Donkey anti-mouse IgG (Yeason). GPCR Extraction Reagent was from Pierce, NP40 lysis buffer was from Beyotime, protease inhibitor mixture cocktail was from Roche Applied Science, rabbit IgG and mouse IgG was from Santa Cruz, and Protein G Dynabeads were from Invitrogen. Tetrodotoxin was from Absin Bioscience. Quinidine was from Macklin, and riluzole was from Meilunbio. All other reagents were purchased from Sigma-Aldrich.

### Molecular cloning

The ion channels used are: Slack-B (Ref Seq: NM_020822.3), Slick (Ref Seq: NM_198503.5), Na_V_1.1 (Ref Seq: NM_001165963.4), Na_V_1.2 (Ref Seq: NM_001040142.2), Na_V_1.3 (Ref Seq: NM_006922.4), Na_V_1.5 (Ref Seq: NM_198056.3), Na_V_1.6 (Ref Seq: NM_014191.4), GluA1 (Ref Seq: XM_032913972.1), GluA2 (Ref Seq: NM_017261.2), GluA3 (Ref Seq: NM_007325.5), and GluA4 (Ref Seq: NM_000829.4). Human Slack-B and Na_V_1.6 were subcloned into the modified pcDNA3.1(+) vector using Gibson assembly. All mutations and chimeras of ion channels were also constructed using Gibson assembly. For GST pull down assay, the segments of Slack and Na_V_1.6 were subcloned into pCDNA3.1(+) and pGEX-4T-1 vector, respectively. For FRET experiments, mVenus-tag was fused to the C-terminus of Na_V_1.6 sequence, and mTFP1-tag was also fused to the C-terminus of Slack sequence.

### Immunoprecipitation

The brain tissues or HEK293T cells co-expressing full-length or fragments of Slack and Na_V_1.6 were homogenized and lysed in GPCR Extraction Reagent (Pierce) with cocktail for 30 min at 4 °C. The homogenate was centrifuged for 20 min at 16 000 g and 4 °C to remove cell debris and then supernatant was incubated with 5 μg Slack antibody (NeuroMab) or Na_v_1.6 antibody (Alomone) for 12 h at 4 °C with constant rotation. 40 μl of protein G Dynabeads (Invitrogen) was then added and the incubation was continued until the next day. Beads were then washed three times with NP40 lysis. Between washes, the beads were collected by DynaMag. The remaining proteins were eluted from the beads by re-suspending the beads in 1×SDS-PAGE loading buffer and incubating for 30 min at 37 °C. The resultant materials from immunoprecipitation or lysates were then subjected to western blot analysis. And the co-immunoprecipitation assays for Slack and Na_V_1.6 were repeated three times independently.

### Western blot analysis

Proteins suspended in 1×SDS-PAGE loading buffer were denatured for 30 min at 37 °C. Then proteins were loaded on 6% or 8% sodium dodecyl sulphate–polyacrylamide gel electrophoresis and transferred onto nitrocellulose filter membrane (PALL). Non-specific binding sites were blocked with Tris-buffered saline-Tween (0.02 M Tris, 0.137 M NaCl and 0.1% Tween 20) containing 5% non-fat dried milk. Subsequently, proteins of interest were probed with primary antibodies for overnight at 4 °C. After incubation with a secondary antibody, immunoreactive bands were visualized using HRP Substrate Peroxide Solution (Millipore) according to the manufacturer’s recommendation.

### GST pull down assay

Plasmids encoding GST-fused Na_V_1.6 segments were transformed into BL21(DE3). After expressing recombinant proteins induced by overnight application of isopropyl-1-thio-β-D-galactopyranoside (IPTG) (0.1mM) at 25℃, the bacteria were collected, lysed and incubated with GSH beads using BeaverBeads GSH kit (Beaver) according to the manufacturer’s instruction.

Plasmids encoding KCNT1 channel or HA-tagged KCNT1 segments were transfected into HEK293T cells using lipofectamine 2000 (Introvigen). 40 h after transfection, cells were lysed in NP40 lysis buffer with inhibitor cocktail, and then centrifugated at 4 ℃, 15000 g for 20 min. The supernatants were incubated with protein-bound beads. The protein-bound beads were washed by washing buffer (50 mM Tris-HCl pH = 7.4, 120 mM NaCl, 2 mM EGTA, 2 mM DTT, 0.1% triton X-100, and 0.2% Tween-20) for 5 min 3 times and then denatured with 1×SDS-PAGE loading buffer and incubating for 30 min at 37 °C. The resultant materials were subjected to western blot analysis.

### Cell culture

The human embryonic kidney cells (HEK293 and HEK293T) were maintained in Dulbecco’s Modified Eagle Medium (DMEM, Gibco) supplemented with 15% Fetal Bovine Serum (FBS, PAN-Biotech) at 37°C and 5% CO_2_. Primary cortical neurons were prepared from either sex of postnatal (P0-P1) homozygous Na_V_1.6 knockout C3HeB/FeJ mice and the wild-type littermate controls. After the mice were decapitated, the cortices were removed and separated from the meninges and surrounding tissue. Tissues were digested in 2 mg/mL Papain (Aladdin) containing 2 μg/mL DNase I (Psaitong) for 30 min followed by centrifugation and resuspension. Subsequently, the cells were plated on poly-D-lysine (0.05 mg/mL) pre-coated glass coverslips in plating medium (DMEM containing 15% FBS), at a density of 4×10^5^ cells/ml, and cultivated at 37°C and 5% CO_2_ in a humidified incubator. 5 h after plating, the medium was replaced with Neurobasal Plus medium (Invitrogen) containing 2% v/v B-27 supplement (Invitrogen), 2 mM Glutamax (Invitrogen), 50 U/mL penicillin and streptomycin (Life Technologies). The primary neurons were grown 6-10 days before electrophysiological recordings with half of the media replaced every three days.

### Voltage-clamp recordings

The plasmids expressing full-length or fragments of Slack and Na_V_ channels (excluding full-length Na_V_1.6) were co-transfected into HEK293 cells using lipofectamine 2000 (Introvigen). To co-express Slack with Na_V_1.6, the plasmid expressing Slack was transfected into a stable HEK293 cell line expressing Na_V_1.6. 18-36 h after transfection, voltage-clamp recordings were obtained using a HEKA EPC-10 patch-clamp amplifier (HEKA Electronic) and PatchMaster software (HEKA Electronic). For all whole-cell patch clamp experiments in HEK cells except for data presented in Fig. S7D, the extracellular recording solution contained (in mM): 140 NaCl, 3 KCl, 1 CaCl_2_, 1 MgCl2, 10 glucose, 10 HEPES, 1 Tetraethylammonium chloride (310 mOsm/L, pH 7.30 with NaOH). The recording pipette intracellular solution (5 mM Na) contained (in mM): 100 K-gluconate, 30 KCl, 15 Choline-Cl, 5 NaCl, 10 glucose,5 EGTA, 10 HEPES (300 mOsm/L, pH 7.30 with KOH). For data presented in Fig. S7D, the extracellular recording solution remains the same as above, the pipette intracellular solution contained (in mM): 140 CsF, 10 NaCl, 5 EGTA, 10 HEPES (pH 7.30 with NaOH), indicating that the unusual right shifts (15∼20 mV) in voltage dependence of Na_V_1.6 were induced by components in pipette solution, not recording system errors. For primary cortical neurons, intracellular solution (0 mM Na) was used to prevent the activation of sodium-activated potassium channels by basal intracellular sodium ions. NaCl in the intracellular solution was replaced with choline chloride in an equimolar concentration. For inside out patch-clamps, the bath solution contained (in mM): 140 NaCl, 1 EDTA, 10 HEPES and 2 MgCl_2_ (310 mOsm/L, pH 7.30 with NaOH). Pipette solution contained (in mM): 130 KCl, 1 EDTA, 10 HEPES and 2 MgCl_2_ (300 mOsm/L, pH 7.30 with KOH). The pipettes were fabricated by a DMZ Universal Electrode puller (Zeitz Instruments) using borosilicate glass, with a resistance of 1.5-3.5 MΩ for whole-cell patch clamp recordings and 8.0-10.0 MΩ for inside-out patch clamp recordings.

All experiments were performed at room temperature. The cells were exposed to the bath solution with quinidine for about one minute before applying voltage protocols. The concentration-response curves were fitted to four-parameter Hill equation:

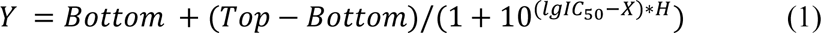

where Y is the value of I_Quinidine_/I_Control_, Top is the maximum response, Bottom is the minimum response, X is the lg of concentration, IC_50_ is the drug concentration producing the half-maximum response, and H is the Hill coefficient. Significance of fitted IC_50_ values compared to control was analyzed using extra sum-of-squares F test.

For data presented in Fig. S7, cells were excluded from analysis if series resistance > 5 MΩ and series resistance compensation was set to 70%∼90%.

The time constants (τ) of activation were fitted with a single exponential equation:

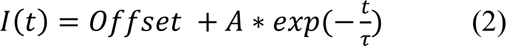

where I is the current amplitude, t is time, offset represents the asymptote of the fit, and A represents the amplitude for the activation or inactivation.

Steady-state fast inactivation (I-V) and conductance-voltage (G-V) relationships were fitting with Boltzmann equations:

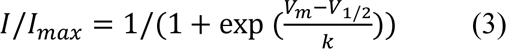

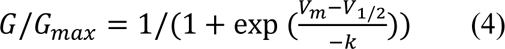

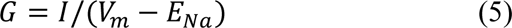

where I is the peak current, G is conductance, V_m_ is the stimulus potential, V_1/2_ is the midpoint voltage, E_Na_ is the equilibrium potential, and k is the slope factor. Significance of fitted V_1/2_ compared to control was analyzed using extra sum-of-squares F test.

Recovery from fast inactivation data were fitted with a single exponential equation:

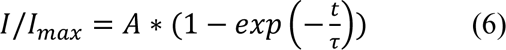

Where I is the peak current of test pulse, I_max_ is the peak current of first pulse, A is the proportional coefficient, t is the delay time between the two pulses, and τ is the time constant of recovery from fast inactivation.

The activation time constants of Na_V_ channels were fitted with a single exponential equation:

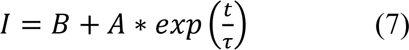

Where I is the current amplitude, A and B are the partition coefficients, t is the time, and τ is the time constant of activation.

### Acute slice preparation and current-clamp recordings

Horizontal slices containing hippocampus were obtained from 6-8 weeks old heterozygous Na_V_1.6 knockout (Scn8a^+/-^) C3HeB/FeJ mice and the wild-type littermate controls. In brief, animals were anesthetized and perfused intracardially with ice-cold modified “cutting solution” containing (in mM): 110 choline chloride, 2.5 KCl, 0.5 CaCl_2_, 7 MgCl_2_, 25 NaHCO_3_, 1.25 NaH_2_PO_4_, 10 glucose; bubbled continuously with 95%O_2_/5%CO_2_ to maintain PH at 7.2. The brain was then removed and submerged in ice-cold “cutting solution”. Next, the brain was cut into 300 μm slices with a vibratome (WPI). Slices were incubated in oxygenated (95% O_2_ and 5% CO_2_) “recording solution” containing (in mM): 125 NaCl, 2.5 KCl, 2 CaCl_2_, 2 MgCl_2_, 25 NaHCO_3_, 1.25 NaH_2_PO_4_, 10 glucose (315 mOsm/L, PH 7.4, 37°C) for 30 min, and stored at room temperature.

Slices were subsequently transferred to a submerged chamber containing “recording solution” maintained at 34-36 °C. Whole-cell recordings were obtained from hippocampal CA1 neurons under a ×60 water-immersion objective of an Olympus BX51WI microscope (Olympus). Pipettes had resistances of 5-8 MΩ. For current-clamp recordings, the external solution (unless otherwise noted) was supplemented with 0.05 mM (2R)-amino-5-phosphonovaleric acid (APV), 0.01 mM 6-cyano-7-nitro-quinoxaline-2,3-dione, 0.01 mM bicuculline and 0.001 mM CGP 55845, and internal pipette solution containing (in mM): 118 KMeSO_4_, 15 KCl, 10 HEPES, 2 MgCl_2_, 0.2 EGTA, and 4 Na_2_ATP, 0.3 Tris-GTP, 14 Tris-phosphocreatinin (pH 7.3 with KOH).

In our whole-cell current-clamp recordings with an Axon 700B amplifier (Molecular Devices), we initially applied a 100 ms, 20 pA test pulse to the recording neurons right after breaking into the whole-cell configuration. A fast-rising component and a slow-rising component of voltage response were clearly visible. Then we zoomed into the fast-rising component of the voltage responses and turned up the pipette capacitance neutralization slowly to shorten the rise time of the fast-rising component until the oscillations of voltage responses appeared. Subsequently, we decreased the capacitance compensation just until the oscillations disappeared. For bridge balance of current-clamp recordings, we increased the value of bridge balance slowly until the fast component of the voltage response disappeared, and the slow-rising component appeared to rise directly from the baseline. Series resistance and pipette capacitance were compensated using the bridge balance and pipette capacitance neutralization options in the Multiclamp 700B command software (Molecular Devices). The bridge balance value was between 20 and 30 MΩ and the pipette capacitance neutralization value was between 3 and 5 pF. For in vitro experiments, the cells were selected by criteria based on hippocampal CA1 cell morphology and electrophysiological properties in the slices. Electrophysiological recordings were made using a Multiclamp 700B amplifier (Molecular Devices). Recordings were filtered at 10 kHz and sampled at 50 kHz. Data were acquired and analyzed using pClamp10.0 (Molecular Devices). Series resistance was in the order of 10–30 MΩ and was approximately 60–80% compensated. Recordings were discarded if the series resistance increased by more than 20% during the time course of the recordings.

### Fluorescence Imaging and FRET Quantification

The spectroscopic imaging was built upon a Nikon TE2000-U microscope. The excitation light was generated by an Ar laser. The fluorescent protein mVenus fused to Na_V_1.6 and mTFP1 fused to Slack were excited by laser line at 500 and 400-440 nm, respectively. The duration of light exposure was controlled by a computer-driven mechanical shutter (Uniblitz). A spectrograph (Acton SpectraPro 2150i) was used in conjunction with a charge coupled device (CCD) camera (Roper Cascade 128B). In this recording mode two filter cubes (Chroma) were used to collect spectroscopic images from each cell (excitation, dichroic): cube I, D436/20, 455dclp; cube II, HQ500/20, Q515lp. No emission filter was used in these cubes. Under the experimental conditions, auto fluorescence from untransfected cells was negligible. Fluorescence imaging and analysis were done using the MetaMorph software (Universal Imaging). User-designed macros were used for automatic collection of the bright field cell image, the fluorescence cell image, and the spectroscopic image. Emission spectra were collected from the plasma membrane of the cell by positioning the spectrograph slit across a cell and recording the fluorescence intensity at the position corresponding to the membrane region (Fig. 2e, dotted lines in red); the same slit position applied to both the spectrum taken with the mTFP1 excitation and the spectrum taken with the mVenus excitation. Using this approach, the spectral and positional information are well preserved, thus allowing reliable quantification of FRET efficiency specifically from the cell membrane. Spectra were corrected for background light, which was estimated from the blank region of the same image.

FRET data was quantified in two ways. First, the FRET ratio was calculated from the increase in mVenus emission due to energy transfer as described in the previous study.^60^ Briefly, mTFP1 emission was separated from mVenus emission by fitting of standard spectra acquired from cells expressiring only mVenus or mTFP1. The fraction of mVenus-tagged molecules that are associated with mTFP1-tagged molecules, Ab, is calculated as

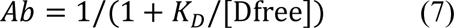

where *K*_D_ is the dissociation constant and [Dfree] is the concentration of free donor molecules. Note that

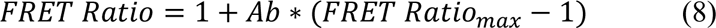

Regression analysis was used to estimate Ab in individual cells. From each cell, the FRET ratio_exp_ was experimentally determined. The predicted Ab value was then computed by adjusting two parameters, FRET Ratio_max_ and apparent *K*_D_. Ab was in turn used to give a predicted FRET ratiopredicted. By minimizing the squared errors (FRET ratio_exp_ – FRET ratio_predicted_)^2^, *K*_D_ was determined.

Second, apparent FRET efficiency was also calculated from the enhancement of mVenus fluorescence emission due to energy transfer^60–63^ using a method as previously described.^64^ Briefly, Ratio A_0_ and Ratio A were measured to calculate FRET efficiency. Ratio A_0_ represents the ratio between tetramethylrhodamine maleimide emission intensities (in the absence of fluorescein maleimide) upon excitation at the donor and acceptor excitation wavelengths^63–65^, and was calculated in the present study at the mVenus peak emission wavelength. A particular advantage of quantifying Ratio A_0_ for FRET measurement is that changes in fluorescence intensity caused by many experimental factors can be cancelled out by the ratiometric measurement. A similar ratio, termed Ratio A, was determined in the presence of mTFP1 in the same way as Ratio A_0_. If FRET occurred, the Ratio A value should be higher than Ratio A_0_; the difference between Ratio A and Ratio A_0_ was directly proportional to the FRET efficiency by the factor of extinction coefficient ratio of mTFP1 and mVenus.^63–65^

### Immunostaining

After deep anesthesia with sodium pentobarbital, mice were sacrificed by perfusion with 0.5% paraformaldehyde and 0.5% sucrose (wt/vol) in 0.1 M phosphate buffer (pH 7.4). The brain was removed and post-fixed in the same fixative for 2 h, and subsequently immersed in 30% sucrose in 0.1 M phosphate buffer for 48 h. Cryostat coronal sections (20 μm) were obtained using a freezing microtome (Leica). The sections were rinsed in 0.01 M phosphate-buffered saline (PBS, pH 7.4), permeabilized in 0.5% Triton X-100 in PBS for 30 min, and incubated in a blocking solution (5% BSA, 0.1% Triton X-100 in PBS, vol/vol) at 20–25 °C for 2 h, followed by overnight incubation at 4 °C with primary antibody to AnkG (1:100, Santa Cruz, sc-31778), Slack (1:100, NeuroMab, 73-051), Na_V_1.2 (1:200, Alomone, ASC-002), Na_V_1.6 (1:200, Alomone, ASC-006), Flag (1:500, Abbkine, ABT2010), and HA (1:500, Abbkine, ABT2040) in blocking solution. After a complete wash in PBS, the sections were incubated in Alexa 488-conjugated doney anti-rabbit IgG and Alexa 594 donkey anti-mouse IgG in blocking solution at 20–25 °C for 2 h. The sections were subsequently washed and rinsed in Dapi solution. Images were taken in the linear range of the photomultiplier with a laser scanning confocal microscope (ZEISS LSM 510 META NLO).

### Adeno-associated virus construction and injection

The adeno-associated viruses (AAVs) and the negative GFP control were from Shanghai GeneChem. Co., Ltd. The full-length Slack_G269S_ sequence (1-1238aa) was ligated into modified CV232 (CAG-MCS-HA-Poly A) adeno-associated viral vector. The Slack’s C-terminus sequence (326-1238aa) and the negative control were ligated into GV634 (CAG-MCS-3×Flag-T2A-EGFP-SV40-Poly A) adeno-associated viral vector. The viruses (>10^11^ TU/ml) were used in the present study.

For dorsal CA1 viral injection, C57BL/6J mice aged 3 weeks were anesthetized with isoflurane and placed in a stereotaxic apparatus (RWD Life Science Co., Ltd.). Using a 5 μL micro syringe (Hamilton) with a 30 gauge needle (RWD Life Science Co., Ltd.), 600 nL of the viruses was delivered at 10 nL/min by a micro-syringe pump (RWD Life Science Co., Ltd.) at the following site in each of the bilateral CA1 regions, using the stereotaxic coordinates: 2.5 mm (anterior-posterior) from bregma, 2 mm (medio-lateral), ± 1.5 mm (dorsal-ventral)^66^. The syringe was left in place for 5 min after each injection and withdrawn slowly. The exposed skin was closed by surgical sutures and returned to home cage for recovery. All the experiments were conducted after at least 3 weeks of recovery. All the mice were sacrificed after experiments to confirm the injection sites and the viral trans-infection effects by checking EGFP under a fluorescence microscope (ZEISS LSM 510 META NLO).

### Kainic acid-induced status epilepticus

KA (Sigma-Aldrich) was intraperitoneally administered to produce seizures with stage IV or higher. The dose of kainic acid used was 28 mg/kg for mice (6–8 weeks)^67,68^. To assess epilepsy susceptibility, seizures were rated using a modified Racine, Pinal, and Rovner scale^45,69^: (1) Facial movements; (2) head nodding; (3) forelimb clonus; (4) dorsal extension (rearing); (5) Loss of balance and falling; (6) Repeated rearing and failing; (7) Violent jumping and running; (8) Stage 7 with periods of tonus; (9) Dead. Seizures was terminated 2 h after onset with the use of sodium pentobarbital (30 mg/kg; Sigma-Aldrich).

### Statistical analysis

For in vitro experiments, the cells were evenly suspended and then randomly distributed in each well tested. For in vivo experiments, the animals were distributed into various treatment groups randomly. Statistical analyses were performed using GraphPad Prism 9 (GraphPad Software) and SPSS 26.0 software (SPSS Inc.). Before statistical analysis, variation within each group of data and the assumptions of the tests were checked. Comparisons between two independent groups were made using unpaired Student’s two-tailed t test. Comparisons among nonlinear fitted values were made using extra sum-of-squares F test. Comparisons among three or more groups were made using one- or two-way analysis of variance followed by Bonferroni’s post hoc test. No statistical methods were used to predetermine sample sizes but our sample sizes are similar to those reported previously in the field^37,70^. All experiments and analysis of data were performed in a blinded manner by investigators who were unaware of the genotype or manipulation. * *p* < 0.05, ** *p* < 0.01, *** *p* < 0.001, **** *p* < 0.0001. All data are presented as mean ± SEM.

## AUTHOR CONTRIBUTIONS

T.Y., Y.J., and H.Y. performed and analyzed voltage-clamp recordings. Y.W. performed and analyzed Western blotting, immunoprecipitation. S.X., C.P., and G.D. performed and analyzed pull down assay. Q.C. performed the immunostaining. H.Z. and F.Y. performed and analyzed the FRET imaging. H.S., N.L., and X.M. performed and analyzed the current clamp recordings. T.Y., H.Y., Z.G., and J.D. performed the molecular cloning. Z.H., T.Y., and Y.W. designed the experiments. Z.H., T.Y., and Y.J. wrote the manuscript. Z.H., F.Y., Y.Y., and Q.S. reviewed the manuscript.

## ACKNOWLEDGMENTS

This work was supported by Chinese National Programs for Brain Science and Brain-like intelligence technology No.2021ZD0202102 to Z.H.; National Natural Science Foundation of China Grant (Nos. 31871083 and 81371432 to Z.H.; Nos. 32000674 to G.D.). We also thank Professor Yousheng Shu at Fudan University for providing the Na_V_1.6-knockout mice.

## DATA AVAILABILITY

The data that support the findings of this study are available from the corresponding author upon reasonable request.

