## Supplementary Figure 1-11 and table 1-4 for "Coupling of Slack and Na_V_1.6 sensitizes Slack to quinidine blockade and guides anti-seizure strategy development"

Tian Yuan *et al.*

**This PDF file includes:**

Fig. S1 to S8

Table S1 to S4

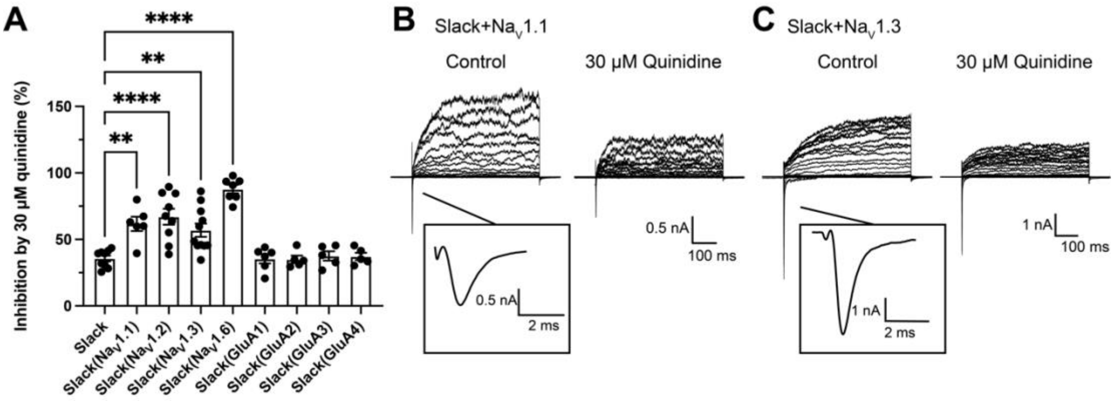


**Figure S1. The sensitivity of Slack to quinidine blockade upon expression of Slack alone and co-expression of Slack with sodium-permeable channels.** (**A**) The inhibitory effects of 30 μM quinidine on Slack upon expression of Slack alone (n = 9) and co-expression of Slack with Na$_V$1.1 (n = 6), Na$_V$1.2 (n = 9), Na$_V$1.3 (n = 11), Na$_V$1.6 (n = 7), GluA1 (n = 6), GluA2 (n = 5), GluA3 (n = 5), or GluA4 (n = 5). ** $p < 0.01$, **** $p < 0.0001$; one-way ANOVA followed by Bonferroni's post hoc test. Upon co-expression of Slack with Na$_V$ subunits, Na$_V$ currents and Slack currents were evoked by applying 600-ms step pulses to voltages varying from -120 mV to +100 mV in 10 mV increments, with a holding potential of -90 mV and a stimulus frequency of 0.20 Hz (pulse protocol 1). Upon co-expression of Slack with AMPAR subunits, the AMPAR currents were elicited by 1 mM Glutamate in the presence of 5 μM Cyclothiazide (CTZ), applied for 10 s before the co-application of CTZ plus Glutamate. After recording AMPAR currents, Slack currents were elicited by a 600-ms step pulse to +100 mV from a holding potential of -90 mV in the absence or presence of 30 μM quinidine in the bath solution (pulse protocol 2). For Slack expressed alone, both pulse protocols were used (n = 5 for pulse protocol 1 and n = 4 for pulse protocol 2). (**B-C**) Example current traces from HEK293 cells co-expressing Slack with Na$_V$1.1 (**B**), Na$_V$1.3 (**C**), before and after application of 30 μM quinidine.

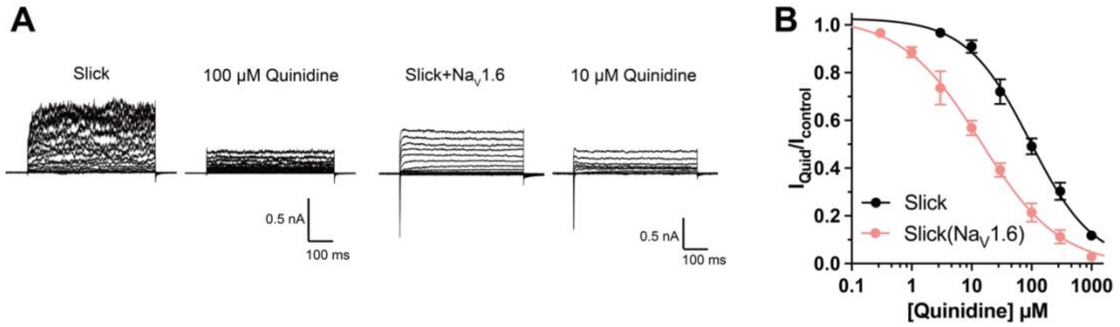


**Figure S2. The sensitivity of Slick to quinidine blockade upon expression of Slick alone or co-expression of Slick with $Na_V1.6$.** (**A**) Example current traces from HEK293 cells expressing Slick alone and co-expressing Slick with $Na_V1.6$. The left traces show the family of control currents; the right traces show the delayed outward current remaining after application of the indicated concentrations of quinidine. (**B**) The concentration-response curves for blocking of Slick by quinidine upon expression of Slick alone (n = 5) and co-expression of Slick with $Na_V1.6$ (n = 7).

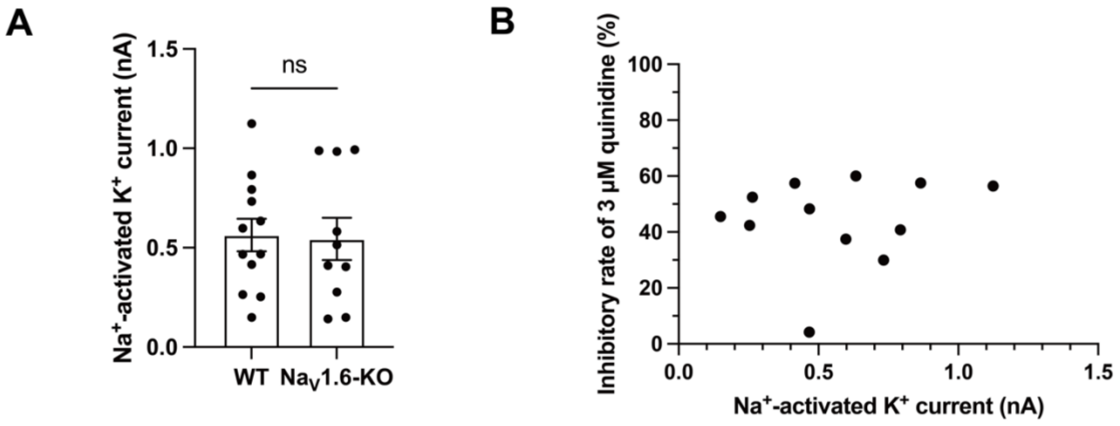


**Figure S3. The amplitudes and sensitivity to quinidine of sodium-activated potassium currents in primary cortical neurons.** (**A**) The amplitudes of $I_{KNa}$ in WT (n=12) and $Na_V1.6$-KO (n=10) neurons. ns, $p > 0.05$, unpaired two-tailed Student's t test. (**B**) The correlation between the amplitudes of $I_{KNa}$ in WT neurons before bath-application of 3 μM quinidine and the inhibitory effect of 3 μM quinidine on $I_{KNa}$ in WT neurons. $r = 0.1555$, $p = 0.6294$, Pearson correlation analysis.

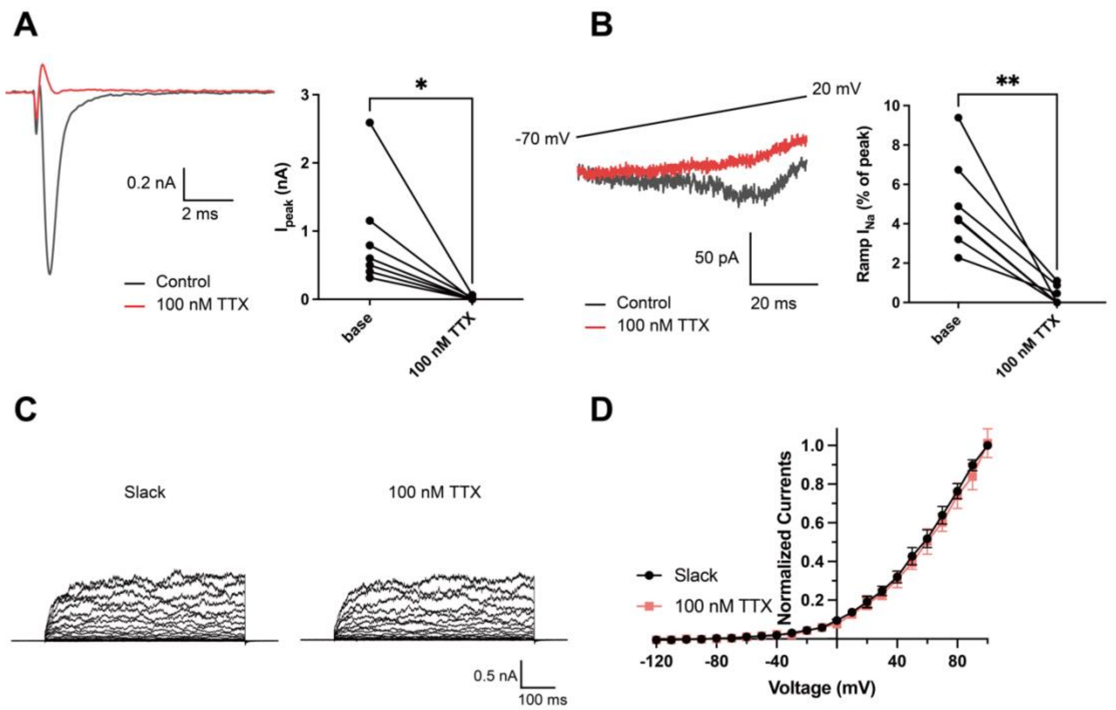


**Figure S4. Effects of 100 nM TTX on Na$_V$1.6 and Slack currents.** (**A**) Example peak currents ($I_{peak}$) at 0 mV of Na$_V$1.6 before (black) and after (red) application of 100 nM TTX in the bath solution. (**B**) Example ramp sodium current (ramp $I_{Na}$) traces evoked during a slow ramp stimulus beginning at −120 mV and ending at 20 mV over a duration of 200 ms (traces from -70 mV to 20 mV were shown). Summarized ramp $I_{Na}$ relative to $I_{peak}$ of Na$_V$1.6 before and after application of 100 nM TTX in the bath solution were shown on the right panel. (**C**) Example current traces of Slack expressed alone evoked from a holding potential of -90 mV. (**D**) I-V relationships of Slack before (black) and after (red) application of 100 nM TTX in the bath solution. * $p < 0.05$, ** $p < 0.01$, paired two-tailed Student's *t* test.

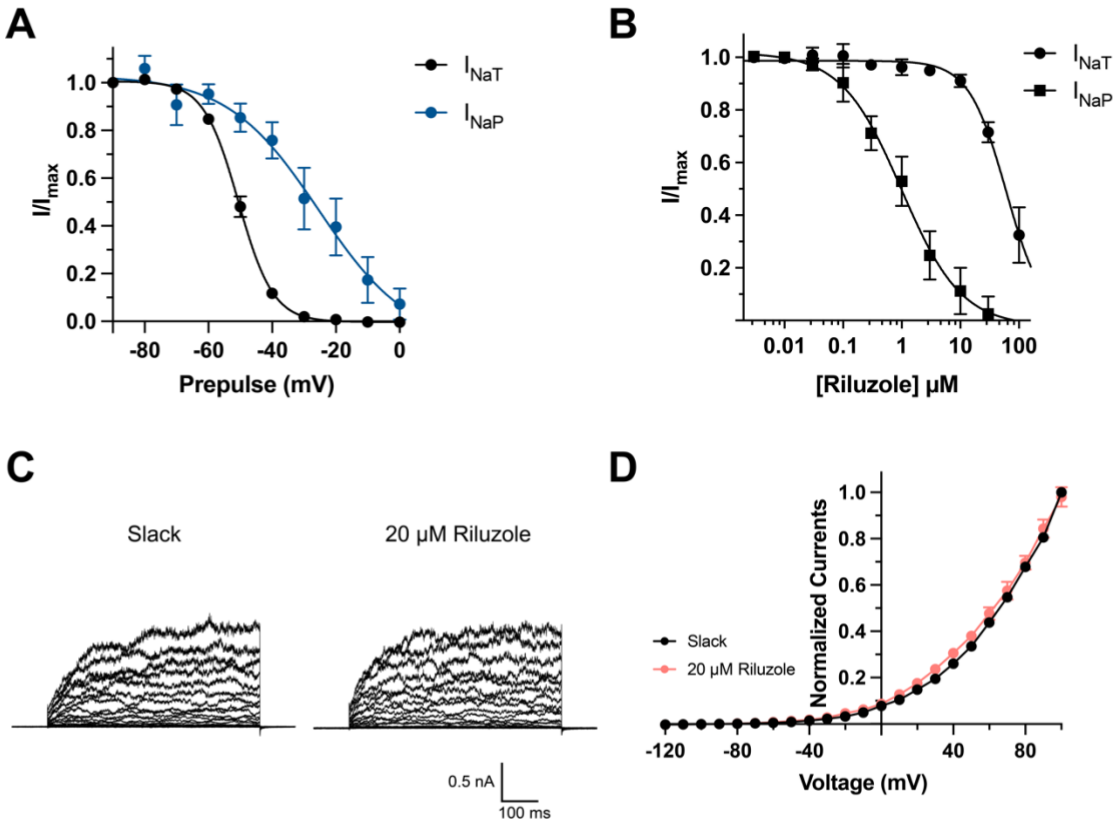


**Figure S5. Effects of depolarized prepulse potentials and riluzole on channels.** (**A**) Inactivation relationships of transient ($I_{NaT}$) and persistent sodium currents ($I_{NaP}$) of $Na_V$1.6 in HEK293 cells. The currents were elicited by a 600-ms step test pulse to 0 mV from a 100-ms prepulse of voltage varying from -90 mV to 0 mV in 10 mV increments. The $I_{NaT}$ represented the peak of sodium current, and the $I_{NaP}$ was assessed at 150 ms in test pulse. (**B**) The concentration-response curves for inhibition of transient (n = 6) and persistent $Na_V$1.6 currents (n = 6) by riluzole upon expression of $Na_V$1.6 alone in HEK293 cells. (**C**) Example current traces from HEK293 cells expressing Slack before and after application of 20 μM riluzole into the bath solution. (**D**) I-V relationships of Slack currents before and after application of 20 μM riluzole (n = 5).

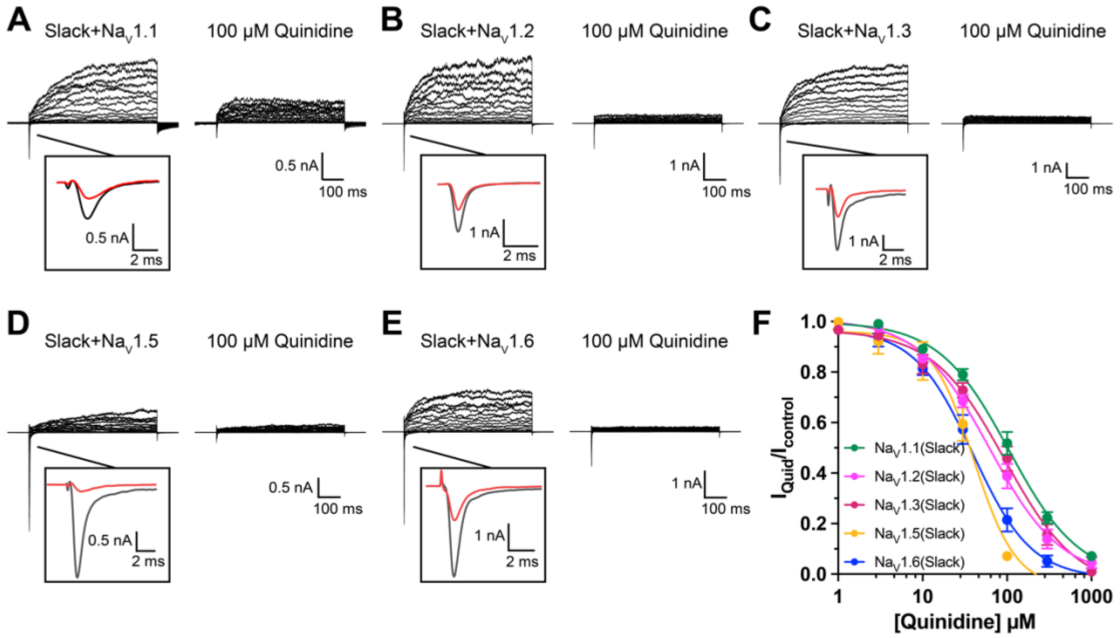


**Figure S6. The sensitivity of Na$_V$ channel subtypes to quinidine blockade.** (**A-E**) Example current traces of Na$_V$1.1 (**A**), Na$_V$1.2 (**B**), Na$_V$1.3 (**C**), Na$_V$1.5 (**D**), and Na$_V$1.6 (**E**) co-expressed with Slack in HEK293 cells. (**F**) The concentration-response curves for blocking of Na$_V$ by quinidine upon co-expression of Na$_V$ with Slack (n = 6 for Na$_V$1.1, n = 3 for Na$_V$1.2, n = 12 for Na$_V$1.3, n = 9 for Na$_V$1.5, and n = 5 for Na$_V$1.6). Please refer to Supplementary Table 2 for $IC_{50}$ values.

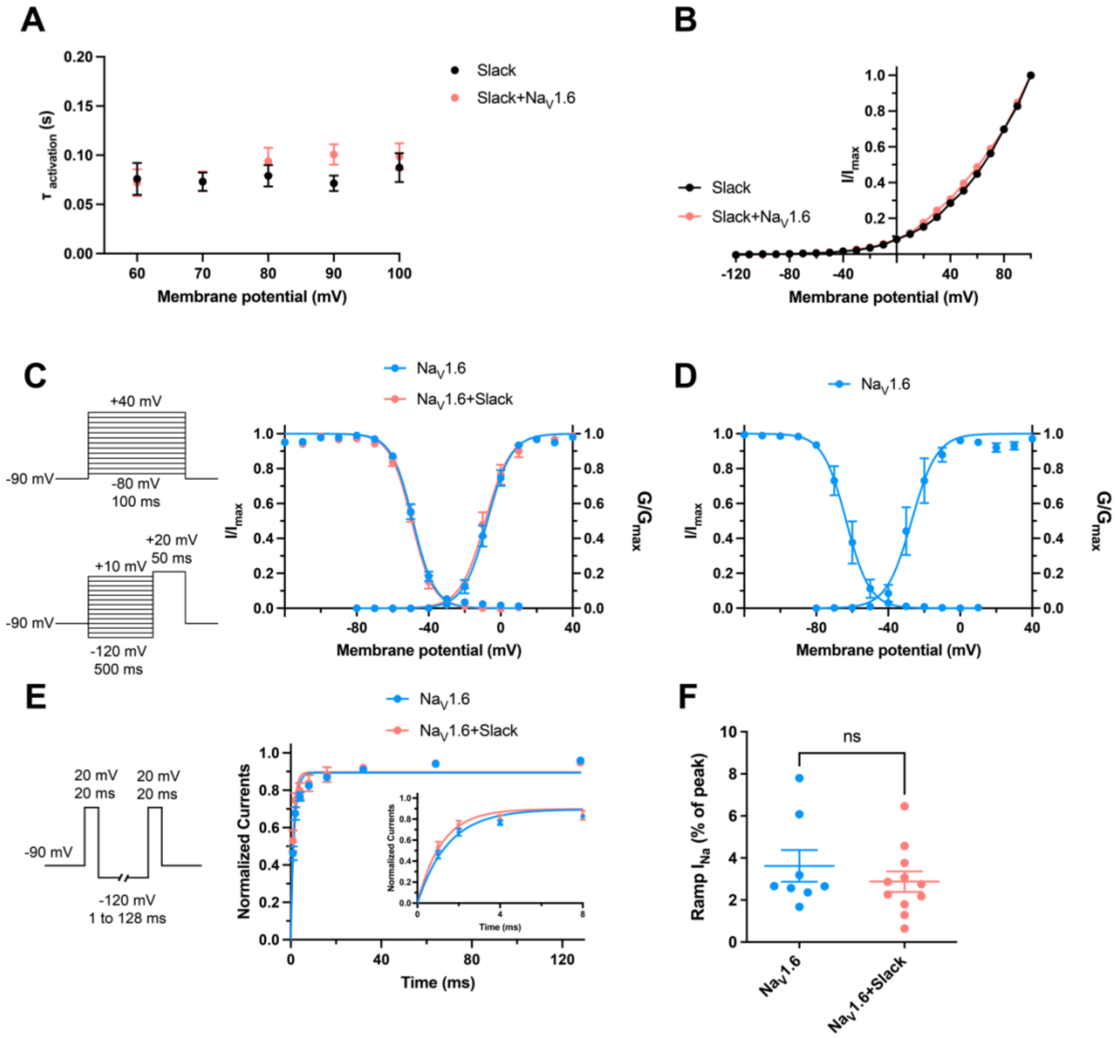


**Figure S7. The dynamic properties of Slack and Nav1.6 channels.** (**A**) Average time constants calculated from single exponential decay fits of activation of Slack currents upon expression of Slack alone and co-expression of Slack with Nav1.6 (n = 10). (**B**) I-V curves of Slack expressed alone (black) or Slack upon co-expressed with Nav1.6 (red). (**C**) Voltage dependence of steady-state activation and steady-state fast inactivation of Nav1.6 sodium currents upon expression of Nav1.6 alone (n = 10 for activation and n = 11 for inactivation; blue) and co-expression of Nav1.6 with Slack (n = 6 for activation and n = 7 for inactivation; red) in HEK293 cells. The pipette solution contained (in mM) 100 K-gluconate, 30 KCl, 15 Choline-Cl, 5 NaCl, 10 glucose,5 EGTA, 10 HEPES. To measure steady-state activation, HEK293 cells were stimulated by a 100-ms step pulses to voltages varying from -80 mV to 40 mV in 10 mV increments, with a holding potential of -90 mV and a stimulus frequency of 0.2 Hz. To measure steady-state fast inactivation, HEK293 cells were stimulated by a 100-ms prepulse to voltages varying from -120 mV to -20 mV in 10 mV increments followed by a test step to -20 mV, with a holding potential of -90 mV and a stimulus frequency of 0.2 Hz. (**D**) Voltage dependence of steady-state activation and steady-state fast inactivation of $Na_V1.6$ sodium currents upon expression of $Na_V1.6$ alone in HEK293 cells (n = 4 for activation and n = 5 for inactivation; using the same voltage protocols as in Fig. S6c). The pipette solution contained (in mM) 140 CsF, 10 NaCl, 1 EGTA, 10 HEPES. (**E**) The voltage protocols and time course for recovery from fast inactivation of $Na_V1.6$ sodium currents upon expression of $Na_V1.6$ (n = 8, blue) and co-expression of $Na_V1.6$ with Slack (n = 7, red). HEK293 cells were stimulated by two 20-ms pulses (prepulse and test pulse) to 20 mV from a recovery potential of -120 mV with recovery phase intervals varying from 1 ms to 128 ms. The solid curves were fitted to the single exponential equation, with a time constant $\tau$ of 1.49 ms for $Na_V1.6$ and a time constant $\tau$ of 1.17 ms for $Na_V1.6$ with Slack. * $p < 0.05$; extra sum-of-squares F test. (**F**) Summarized ramp $I_{Na}$ relative to $I_{peak}$ of $Na_V1.6$ upon expression of $Na_V1.6$ alone (n = 8) and co-expression of $Na_V1.6$ with Slack (n = 11) (using the same protocols as in Fig. S3b). Please refer to Supplementary Table 3 for values of half-maximal activation and inactivation ($V_{1/2}$) and slope factors (k).

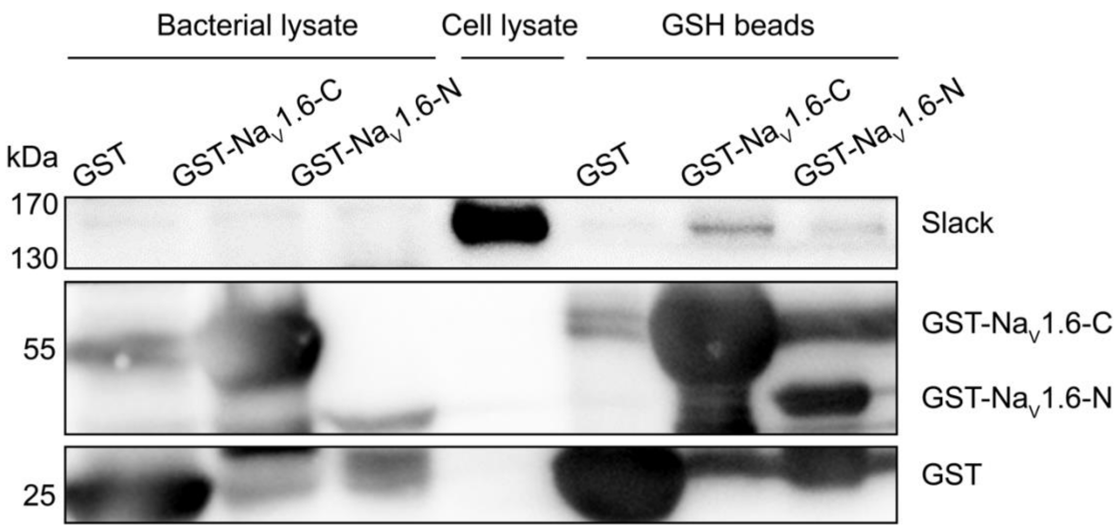


**Figure S8. GST pull down assay of Slack with the N- and C-termini of Na$_V$1.6.** The GST-fused Na$_V$1.6's termini were separately expressed in BL21(DE3) and captured by GSH-beads. The GST-fused proteins were subsequently incubated with cells lysates of HEK293T cells expressing Slack. Input (Bacterial lysate) volume corresponds to 20% of the total lysates for pull down.

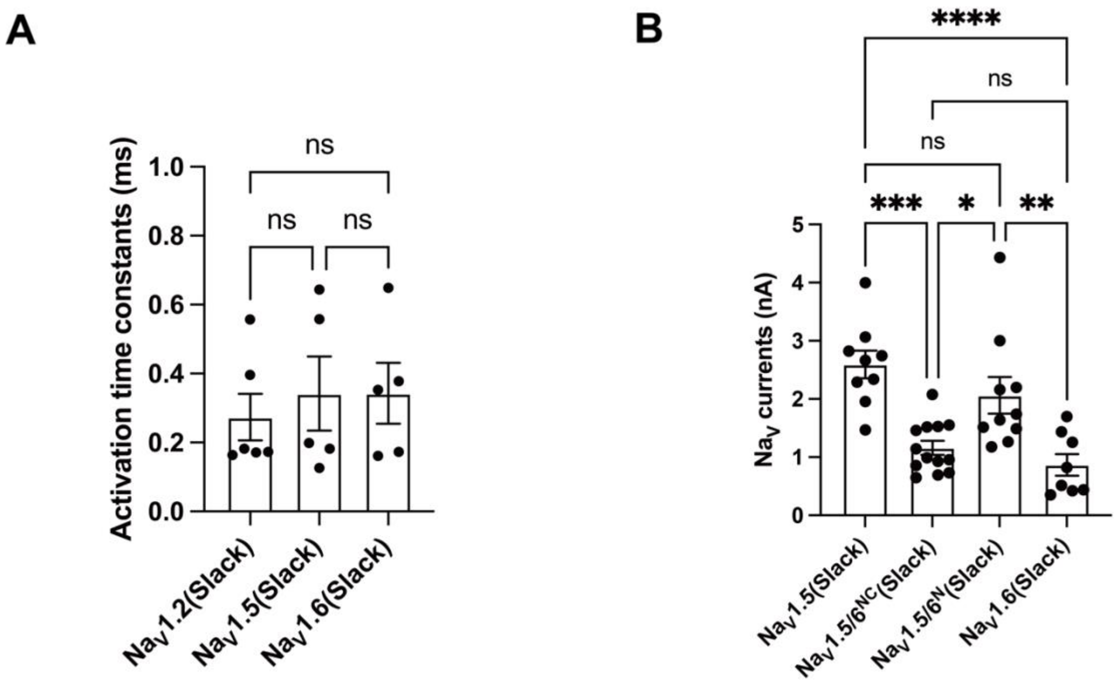


**Figure S9. Comparison of the activation and amplitudes of Na$_V$ channel subtypes currents upon co-expressed with Slack.** (**A**) The activation time constants of peak sodium currents in HEK293 cells co-expressing Na$_V$1.2 (n=6), Na$_V$1.5 (n=5), and Na$_V$1.6 (n=5) with Slack, respectively. ns, $p > 0.05$, one-way ANOVA followed by Bonferroni's post hoc test. (**B**) Comparison of peak sodium current amplitudes of Na$_V$1.5 (n=9), Na$_V$1.5/6$^{NC}$ (n=13), NaV1.5/6$^{N}$ (n=10), and Na$_V$1.6 (n=8) upon co-expressed with Slack in HEK293 cells. ns, $p > 0.05$, * $p < 0.05$, ** $p < 0.01$, *** $p < 0.001$, **** $p < 0.0001$; one-way ANOVA followed by Bonferroni's post hoc test.

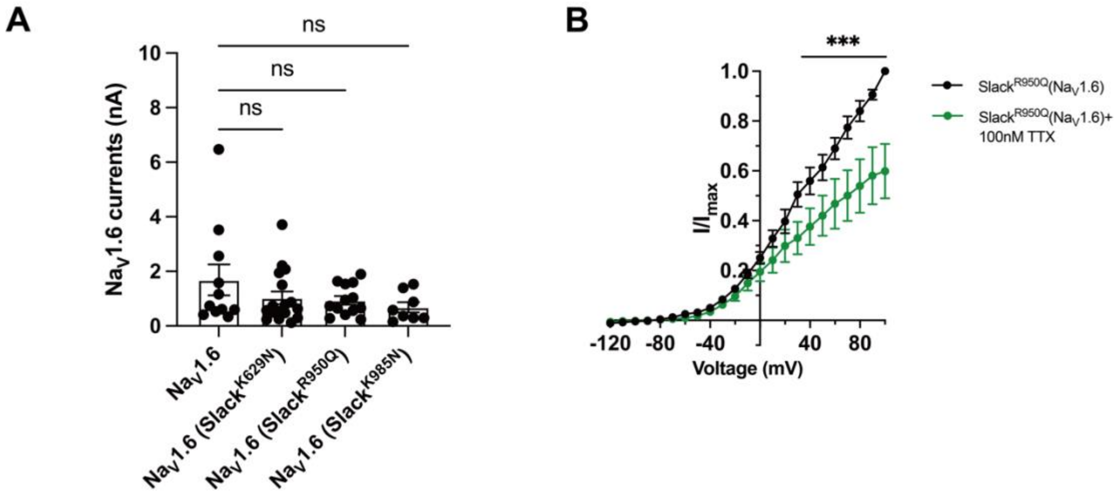


**Figure S10. The $Na^{+}$-mediated currents coupling of Slack mutant variants and $Na_V1.6$ upon co-expression in HEK293 cells.** (**A**) Comparison of $Na_V1.6$ sodium current amplitudes upon expression of $Na_V1.6$ alone (n=11) and co-expression of $Na_V1.6$ with epilepsy-related Slack mutant variants [$Slack^{K629N}$ (n=17), $Slack^{R950Q}$ (n=13), and $Slack^{K985N}$ (n=8)]. ns, $p > 0.05$, one-way ANOVA followed by Bonferroni's post hoc test. (**B**) The current amplitudes of $Slack^{R950Q}$ before (black) and after (green) bath-application of 100 nM TTX upon co-expression with $Na_V1.6$ in HEK293 cells (n=5). ***$p < 0.001$, Two-way repeated measures ANOVA followed by Bonferroni's post hoc test.

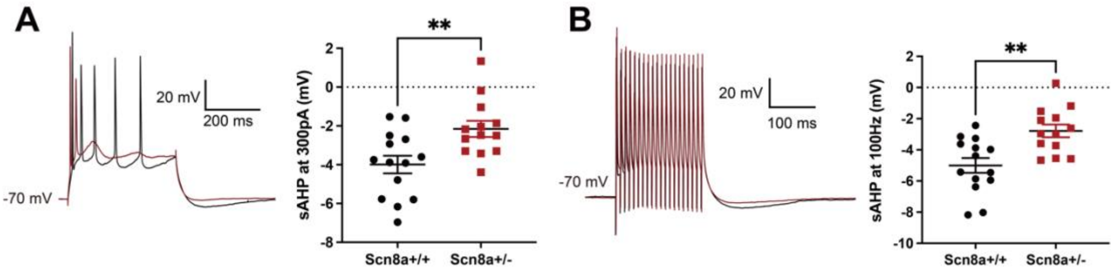


**Figure S11. Heterozygous knockout of $Na_V$1.6 decreases the amplitude of slow AHP in hippocampal CA1 pyramidal neurons.** (**A**) Action potentials elicited at 300 pA followed by a slow AHP in wild-type (WT) neurons (n = 14, black) and heterozygous $Na_V$1.6 knockout (Scn8a$^{+/-}$) neurons (n = 13, red). The amplitude of slow AHP reduced in the Scn8a$^{+/-}$ neurons. ** $p < 0.01$; unpaired two-tailed Student's *t* test. (**B**) Action potentials elicited in the 100 Hz pulse train followed by a slow AHP in WT (n = 14, black) and Scn8a$^{+/-}$ neurons (n = 13, red). The amplitude of slow AHP reduced in the Scn8a$^{+/-}$ neurons. ** $p < 0.01$; unpaired two-tailed Student's *t* test.

**Table S1. The sensitivity of Slack to quinidine blockade upon expression of Slack alone and co-expression of Slack with Na$_V$1.x**

| Channels | IC$_{50}$ (μM) | 95% CI | n |
|---|---|---|---|
| **Slack** | 85.13 | 58.29 to 126.8 | 6 |
| **Slack(Na$_V$1.1)** | 24.83 | 16.36 to 26.55 | 7 |
| **Slack(Na$_V$1.2)** | 14.83 | 7.90 to 28.30 | 10 |
| **Slack(Na$_V$1.3)** | 23.64 | 10.88 to 50.80 | 13 |
| **Slack(Na$_V$1.5)** | 29.46 | 11.58 to 78.67 | 9 |
| **Slack(Na$_V$1.6)** | 0.87 | 0.58 to 1.30 | 19 |

**Table S2. The sensitivity of $Na_V$ channel subtypes to quinidine blockade upon expression of $Na_V$1.x alone and co-expression of $Na_V$1.x with Slack**

| **Channels** | **$IC_{50}$ (μM)** | **95% CI** | **n** |
|---|---|---|---|
| **$Na_V$1.1** | 129.84 | 115.06 to 146.52 | 5 |
| **$Na_V$1.2** | 79.71 | 57.75 to 110.02 | 3 |
| **$Na_V$1.3** | 63.77 | 52.43 to 77.56 | 6 |
| **$Na_V$1.5** | 35.61 | 28.88 to 43.91 | 6 |
| **$Na_V$1.6** | 51.50 | 35.77 to 74.15 | 4 |
| **$Na_V$1.1(Slack)** | 105.72 | 81.10 to 137.81 | 6 |
| **$Na_V$1.2(Slack)** | 62.04 | 49.29 to 78.11 | 3 |
| **$Na_V$1.3(Slack)** | 94.04 | 63.64 to 138.98 | 12 |
| **NaV1.5(Slack)** | 40.23 | 24.18 to 66.94 | 9 |
| **$Na_V$1.6(Slack)** | 39.41 | 28.10 to 55.28 | 5 |

**Table S3. Biophysical characteristics of Nav1.6 expressed alone and Nav1.6 upon co-expression with Slack**

| **Channels** | **Pipette solution** | **Steady-state activation** | | | **Steady-state fast inactivation** | | |
|---|---|---|---|---|---|---|---|
| | | $V_{1/2}$ (mV) with 95%CI | k with 95%CI | n | $V_{1/2}$ (mV) with 95%CI | k with 95%CI | n |
| **Nav1.6** | K-gluconate-based | -7.38 (-8.36 to -6.40) | 6.69 (5.83 to 7.56) | 10 | -48.71 (-49.37 to -48.06) | 6.04 (5.46 to 6.62) | 11 |
| **Nav1.6 (Slack)** | K-gluconate-based | -8.52 (-9.63 to -7.41) | 7.06 (6.08 to 8.04) | 6 | -49.48 (-50.34 to -48.62) | 6.07 (5.30 to 6.83) | 7 |
| **Nav1.6** | CsF-based | -27.03 (-29.26 to -24.79) | 7.06 (5.09 to 9.03) | 5 | -63.35 (-64.86 to -61.83) | 6.52 (5.19 to 7.85) | 5 |

**Table S4. The sensitivity of Slack mutant variants to quinidine blockade upon expression of Slack mutant variants alone and co-expression of Slack mutant variants with Na$_V$1.6**

| Channels | IC$_{50}$ (μM) | 95%CI | n |
|---|---|---|---|
| **Slack$^{K629N}$** | 19.01 | 12.50 to 28.90 | 8 |
| **Slack$^{K629N}$(Nav1.6)** | 0.26 | 0.09 to 0.73 | 8 |
| **Slack$^{R950Q}$** | 25.06 | 13.12 to 47.89 | 7 |
| **Slack$^{R950Q}$(Nav1.6)** | 0.34 | 0.13 to 0.85 | 5 |
| **Slack$^{K985N}$** | 46.10 | 31.52 to 67.42 | 5 |
| **Slack$^{K985N}$(Nav1.6)** | 2.41 | 1.52 to 3.80 | 7 |
